# Dual Traditional Chinese Medicine-Preconditioned Stem Cell Secretomes in Coaxial Electrospun Nanofibers Synergistically Accelerate Diabetic Wound Regeneration

**DOI:** 10.64898/2026.08.25.746942

**Authors:** Kuo-Hui Chiu, Lin-Chu Huang, Wen-Ling Wang, Yi-Hui Lai, Chun-Hsu Yao

## Abstract

Chronic diabetic wounds resist healing due to impaired angiogenesis, stalled cellular migration, and persistent inflammation, a challenge further compounded by the rapid degradation of therapeutic growth factors in the proteolytic wound bed. To overcome this, Traditional Chinese Medicine (TCM) compounds are employed not as standalone drugs, but as biomolecular stimuli to precondition the secretome of Wharton’s Jelly-derived mesenchymal stem cells (WJMSCs). To overcome these critical translational barriers, this study engineers a core-shell coaxial electrospun nanofibrous scaffold (polyvinyl alcohol core/gelatin shell) designed for the stabilizing and sustained dual-delivery of biologics. We introduce a novel synergistic payload with WJMSCs conditioned medium (WJMSCs-CM) uniquely primed by two specific chinese herbal compounds, Astragaloside IV (AS-IV) and Formononetin (FMN). This core-shell architecture provides native-like contact guidance for cells while converting conventional burst release into a sustained, weeks-long elution. *In vitro*, this functionalized scaffold restores Akt/eNOS signaling, rescues cellular viability, and promotes robust tube formation in high-glucose-stressed fibroblasts and endothelial cells. *In vivo*, within an STZ-induced diabetic rat model, the application of this WJMSCs-CM-loaded coaxial scaffold actively inhibits early inflammation and comprehensively accelerates healing, driving near-complete wound closure (98.2 ± 1.5% by day 21), mature collagen deposition, and hair follicle neogenesis. Ultimately, this bio-instructive platform successfully integrates physical structural cues with sustained biochemical signaling, offering a potent, multifaceted strategy for chronic wound regeneration.

**Graphical abstract:** (Figure created in BioRender. Huang, L. C. (2026))

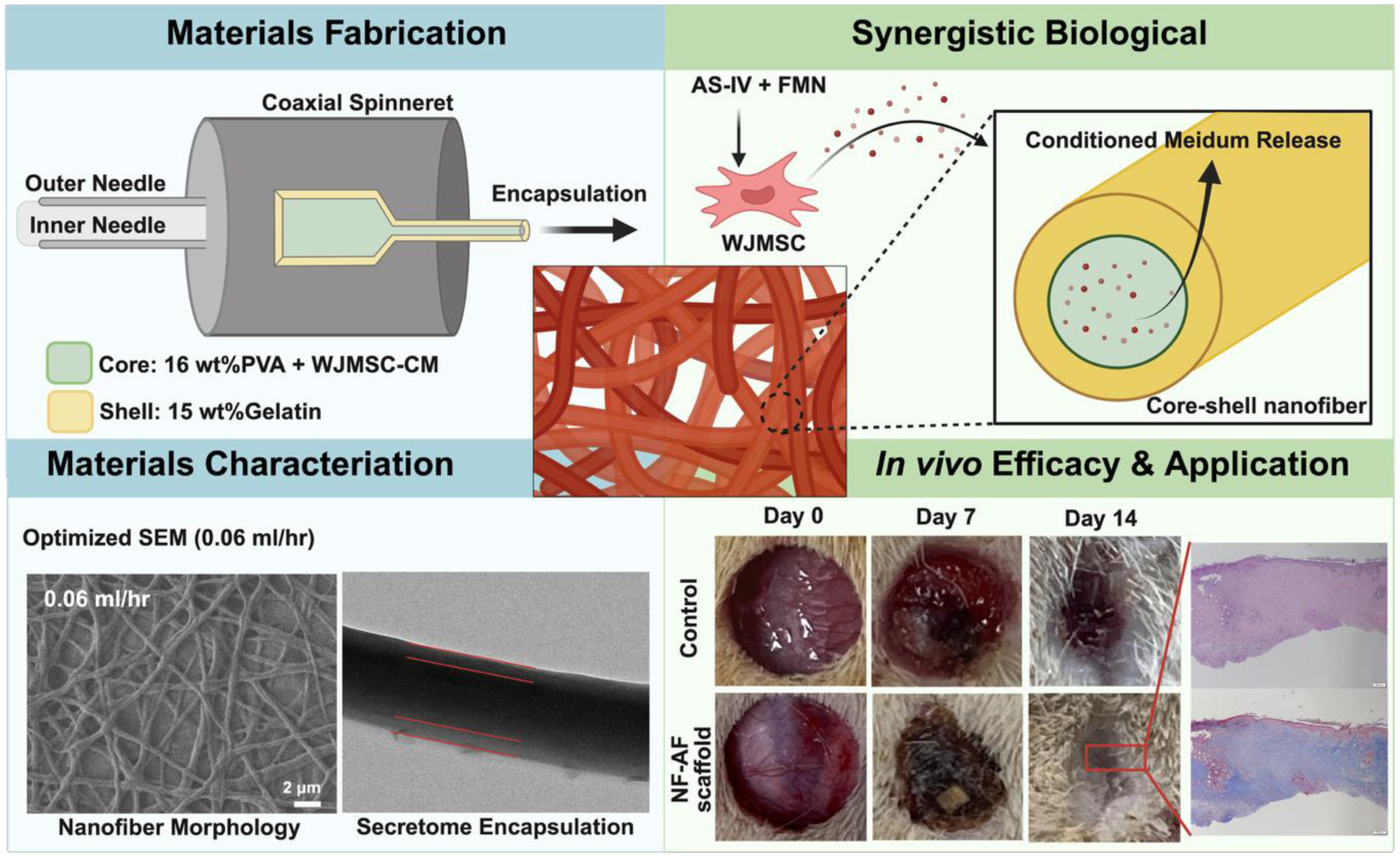

## Introduction

Chronic wounds represent a severe and debilitating complication of diabetes mellitus, frequently culminating in infection, tissue necrosis, and lower-limb amputation^[1]^. The pathology of the diabetic wound microenvironment is driven by a complex breakdown in cellular communication, characterized by impaired angiogenesis, delayed cell proliferation, and a persistent inflammatory state. Current clinical interventions, which predominantly rely on passive dressings to maintain moisture balance or topical antimicrobial agents^[2]^, often fail to actively correct these underlying cellular dysfunctions or restore tissue function. Consequently, there remains a pressing and unmet need for advanced wound care strategies that transcend passive protection to dynamically instruct vascularization and tissue regeneration.

To address these challenges, therapeutic strategies must act synergistically on multiple biological fronts. While the secretome of Wharton’s Jelly-derived Mesenchymal Stem Cells (WJMSCs) has emerged as a potent driver of wound healing, and Traditional Chinese Medicine (TCM) offers a vast reservoir of bioactive compounds known to promote regenerative processes, both face distinct translational hurdles^[3]^. The direct topical application of TCM is often hindered by poor bioavailability and rapid clearance in the wound bed. Conversely, the baseline secretome of native stem cells may lack the highly concentrated specific factors required to overcome the severe ischemia and robust inflammatory cascades unique to diabetic wounds^[4]^.

To address these limitations, the present study proposes a strategy of biomolecular preconditioning, leveraging specific TCM compounds as bioactive stimuli to modulate the WJMSCs secretome. Rather than relying on the basal secretion profile of stem cells, this preconditioning approach employs selected herbal compounds to stimulate the cells in culture, yielding a conditioned medium (CM) with an amplified expression of critical angiogenic and immunomodulatory cytokines^[5]^. The objective of this strategy is to generate a bio-instructive therapeutic cocktail that is significantly more targeted and potent for the diabetic wound microenvironment than either component in isolation. Nevertheless, the successful clinical translation of such protein-rich biological therapies faces a critical delivery hurdle. The diabetic wound bed is characterized by elevated protease activity such as matrix metalloproteinases (MMPs) and inflammatory enzymes, resulting in the rapid degradation of exogenous growth factors^[6]^. Furthermore, conventional topical delivery methods frequently result in an uncontrolled "burst release", which is inadequate for providing the sustained biochemical signaling necessary for comprehensive tissue repair, which can extend over weeks ^[7]^. Therefore, there is an imperative need for an advanced delivery vehicle that possesses the capability to stabilize these labile proteins, thereby dictating a precise release profile and physically guiding cell behavior.

In order to meet these exacting criteria, the present study engineered a core-shell nanofibrous architecture via coaxial electrospinning. Electrospinning has been recognized as a leading technique for fabricating nanofibrous scaffolds that replicate the physical topography of the ECM, thereby providing an optimal template for cell attachment and migration^[8]^. In contrast to conventional monolithic electrospinning, which subjects sensitive cytokines to denaturing organic solvents and exhibits uncontrolled elution, the coaxial design offers two distinct functional advantages^[9,10]^. The advanced core-shell coaxial design facilitates the concurrent extrusion of two distinct fluids: a biodegradable polymer solution that constitutes the protective shell and an aqueous solution containing the bioactive herbal-induced CM within the core. The biodegradable poly(vinyl alcohol) (PVA) core safely encapsulates the TCM-preconditioned CM, while the gelatin shell serves as a physical barrier against proteolytic degradation and precisely regulates molecular diffusion^[11]^. Crucially, the nanoscale topography of the gelatin shell mimics the native extracellular matrix (ECM), thereby providing essential guidance for cellular adhesion and migration ^[12,13]^.

The primary purpose of this study is to apply this WJMSC condition medium (WJMSCs-CM)-loaded coaxial nanofibrous scaffold to comprehensively enhance and accelerate diabetic wound healing. The integration of the physical structural cues of the scaffold with the sustained release of the preconditioned secretome is a specific design feature of this platform, the purpose of which is to actively inhibit persistent inflammation, promote robust tissue regeneration, and drive neo-vascularization in the chronic wound microenvironment.

## Methods and materials

### Material

Type A gelatin derived from porcine skin was acquired from Sigma (St. Louis, MO, USA). Poly(vinyl alcohol) (PVA, degree of polymerization ≈1400; Mw = 60,000–67,000 g/mol; fully hydrolyzed, >99%) was used in this study. was provided by SHOWA (Tokyo, Japan). Astragaloside IV (AS-IV), Dracorhodin perchlorate (DP), Formononetin (FMN), Boswellic acid (BA), Catechin (CA), and Salidroside (SAL) were procured from MedChemExpress (Monmouth Junction, NJ, USA).

### Cell culture

Wharton’s jelly-derived mesenchymal stem cells (WJMSCs, BCRC RM60596, Bioresource Collection and Research Center, Food Industry Research and Development Institute, Hsinchu, Taiwan) and human umbilical vein endothelial cells (HUVECs, BCRC H-UV001) were obtained from the Bioresource Collection and Research Center (BCRC, Taiwan), while human dermal fibroblasts (HDFa) were purchased from Gibco (Cat. No. C-013-5C). All cell types were maintained at 37 °C in a humidified atmosphere containing 5% CO₂, with their respective complete media replenished every 2–3 days. Upon reaching 80%–90% confluence, the cells were washed twice with PBS (Gibco, Grand Island, NY, USA) and incubated with 0.25% Trypsin-EDTA at 37 °C for cell detachment. Once the cells rounded up, the enzymatic reaction was promptly neutralized by adding a double volume of serum-containing complete medium. The detached cells were collected, centrifuged at 1000 rpm for 5 minutes, resuspended in fresh complete medium, and subsequently subcultured at split ratios ranging from 1:2 to 1:4.

### Condition medium collection

To collect the CM, we seeded 2 × 10⁵ WJMSCs/well in 6-well plates, and co-cultured with six different Chinese herbal medicines as listed in materials in different concentration (50, 10, 1, 0.1, 0.01, 0 μg/ml) to stimulate the cell behavior. WJMSCs cells were subjected to herbal treatment for 2 days before collecting CM. On day 3, we first rinsed the cells with PBS and replaced the supernatant with fresh medium. After a 24-hour treatment, the CM was collected and stored at −80 °C prior to lyophilization using a freeze-dryer (Kingmech Co., Ltd., CTD, model FD4.5-12P, serial no. 1517). The lyophilized CM was reconstituted in sterile distilled water to prepare a 10× stock solution, followed by sterilization through a 0.22 µm syringe filter for subsequent experiments.

### Cytokine array

To evaluate the cytokine concentration in the CM, we analyzed the CM with Human Cytokine Antibody Array (membrane-based array containing 80 targets; Abcam, Cambridge, UK; Cat. No. ab133998) following the protocol from manufacturer’s instructions. Briefly, CM samples were incubated overnight at 4 °C with pre-spotted antibody membranes, followed by incubation with a biotin-conjugated detection antibody and horseradish peroxidase (HRP)-linked streptavidin. Chemiluminescent signals were detected and quantified by densitometry using ImageJ software (NIH, Bethesda, MD, USA). Each membrane was normalized to internal positive control spots to account for signal intensity variations.

### Cell viability assay

WJMSCs were seeded into 96-well plates at a density of 5 × 10³ cells/well and employed to evaluate the cytotoxicity of six Chinese herbal medicines. WJMSCs were seeded in a 96-well plate and treated with various concentrations of AS-IV, DP, FMN, BA, CA, and SAL for 2 days at 37 °C in a humidified atmosphere containing 5% CO₂. The cells were then assessed for cell viability by MTT assay. HDFa and HUVECs were also seeded into 96-well plates at a density of 5 × 10³ cells/well and 1 × 10⁴ cells/well, and evaluated the cell viability of CM-induced recovery effect by MTT assay.

### Wound healing assay

We utilized an ibidi culture-insert 2-well system (ibidi, Gräfelfing, Germany) to assess the wound healing behavior. HDFa (1 × 10^4^ cells/0.22 cm^2^) were seeded into each chamber of the insert and cultured until confluence. The 2-well inserts were then removed to create a cell-free gap. Cells were subsequently treated with CM and experimental medium under high-glucose conditions. Cell migration was monitored by inverse microscopy at 0 h and 24 h. The wound closure was imaged and analyzed using ImageJ software according to the following equation:

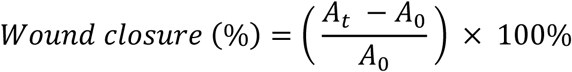

where A_0_ represents the starting wound area at 0 h, and A_t_ denotes the wound area after the treatment.

### Western blotting

The whole cell extract of the HDFa cells was isolated using the lysis buffer. In short, equal amounts of protein were subjected to electrophoresis in SDS-PAGE and transferred onto the polyvinylidene difluoride membranes. The membranes were blocked with 5% non-fat milk in TBST for 1 hour at room temperature, followed by the application of primary antibodies, which were employed for the protein expression analysis: rabbit anti-eNOS [p Ser1177] (Novus Biologicals; 1:2000, Cat# NBP3-05432), rabbit anti-eNOS (Novus Biologicals; 1:2000, Cat# NB300-500), rabbit anti-phospho-Akt (Ser473) (Cell Signaling Technology; 1:2000, Cat# 9271), rabbit anti-AKT (HL2915) (Novus Biologicals; 1:2000, Cat# NBP3-48751), and rabbit anti-GAPDH (Elabscience; 1:5000, Cat# E-AB-20059) were used at 4 °C overnight. Subsequently, membranes were incubated with a goat anti-rabbit IgG H&L (HRP) secondary antibody (Abcam; 1:5000, Cat# ab7090) at room temperature for a minimum of 1 h, and protein bands were detected with an enhanced chemiluminescence (ECL) detection system.

### Tube formation analysis

The µ-Slide angiogenesis (ibidi) is applied for the tube formation assay. Matrigel (10 µl) was coated on the surface of each well and incubated at 37 °C for 30 min. Subsequently, HUVECs cells were suspended in a culture medium with different CM at a density of 1 × 10^4^ cells/well were added to each well after 4 h of cultivation. The tube formation assay was imaged using an Olympus CellSens Dimension microscope system (Olympus Corporation, Tokyo, Japan) and quantified using ImageJ software with the Angiogenesis Analyzer macro toolset (https://imagej.net/ij/macros/toolsets/Angiogenesis%20Analyzer.txt).

### Electrospun nanofibers fabrication

Coaxial electrospun nanofibers featuring a poly(vinyl alcohol) (PVA) core and a gelatin-shell were fabricated using a dual-needle configuration. To prepare the core solution, a 16 wt% PVA solution was dissolved in a mixture of ethanol and deionized water (1:9 v/v) at 60 °C. Concurrently, a condensed (50×) conditioned medium was incorporated into the PVA solution to achieve a final concentration of 5×. For the shell solution, type A gelatin was dissolved at 15 wt% in an equal volume mixture of ethanol and 10× phosphate-buffered saline (PBS) (1:1 v/v) at 40 °C.

The core and shell solutions were loaded into separate 10 mL syringes. Electrospinning was performed using a coaxial spinneret comprising an inner 24G needle and an outer 19G needle. To maintain a 1:1 mass ratio of PVA to gelatin in the resulting composite nanofibers, the feeding flow rates for both the core and shell layers were systematically varied among 0.06, 0.08, 0.1, 0.2, 0.3, and 0.4 mL/h. The electrospinning process was conducted with a spinneret-to-collector distance of 15 cm under an applied high voltage of 20 kV for either 1 or 2 h. Following fabrication, the coaxial nanofibrous membranes were cross-linked via exposure to 50 wt% glutaraldehyde vapors for 45 min, followed by overnight sterilization under ultraviolet (UV) radiation.

### Scanning electron microscopy

Scanning electron microscopy (SEM, JEOL JSM-6700F, Japan) was employed to analyze the dimensions and morphology of the PVA/gelatin core-shell nanofiber at an accelerating voltage of 3 kV. The gold coating was conducted for 90 sec prior to observation. The samples which have at least more than 50 individual fibers in the SEM images were imaged and calculated the mean ± standard deviation of fiber diameter by ImageJ.

### Porosity analysis

The surface porosity of the scaffolds was determined using 2D image analysis. We evaluated the surface area of pores by high-resolution images obtained from the SEM with ImageJ area calculation following the equation,

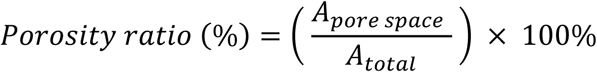

where A_pore space_ represent the area occupied by pores, and Atotal is the total surface area of the scaffold. Threshold was applied to the grayscale images to distinguish the fibers from the void spaces. A minimum of three different regions per sample and minimum of three scaffolds were analyzed to ensure statistical significance.

### Swelling and degradation behavior

The nanofibers were placed in 24-well plates and flattened with an O-ring. One ml of ddH2O was added to each sample and incubated for 0, 3, 6, 12, 24, 48, and 72 h. The swelling ratio of the scaffold was calculated based on the fiber diameter changes with the following equation.

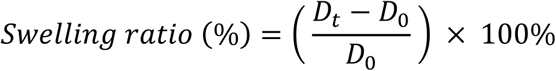

D_0_ is the initial diameter at 0 h, and D_t_ is the diameter at the indicated time point. Triplicate experiments with three individual scaffolds were examined to indicate the statistics.

In addition, to evaluate the degradation behavior, PBS was added to each dry sample, and the samples were then incubated for 7, 14, 21, and 28 days. Each sample was removed and gently blotted with filter paper. Subsequently, the samples were lyophilized and weighed. The weight loss was calculated for the degradation ratio with the following equation,

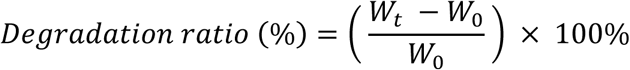

where W_0_ is the initial weight at 0 h in its dry state, and W_t_ is the weight of the specimen after submersion in PBS for the specific time.

### Transmission electron microscopy

The morphology of the coaxial nanofibers was characterized using a transmission electron microscope (TEM; JEOL JEM-F200, Japan) at an acceleration voltage of 120 kV. The coaxially electrospun nanofiber samples for the TEM observation were prepared by directly depositing the as-spun fibers directly deposited onto copper grids for 30 sec during the electrospun process, which had been coated with a supportive formvar film. High-magnification images were captured to clearly resolve the interface between the core and the shell. ImageJ software was employed to quantify the diameters of the core and shell.

### Growth factor release analysis

The coaxial membrane samples were each soaked and pressed by an o-ring, making sure the membrane avoided floating in PBS. In this study, the conditioned medium release assay was conducted in accordance with ISO 10993-12, and the extraction medium volume was determined based on the specified surface area-to-volume ratio. Based on the surface area of the sample, an extraction volume of four hundred microliters was used. Samples were collected on days 1, 3, 5, 7, and 14, and stored at -20 °C for ELISA assay kit analysis.

### Immunofluorescence staining

HDFa (1 × 10^4^ cells/well) was seeded on the membrane in a 24-well plate overnight, and after 1 day, the supernatant was collected for the other experiment. The nanofiber samples were fixed with 4 % formaldehyde for 20 min at room temperature and then washed at least two times with 1XPBS. The samples were permeabilized using 0.1 v/v% Triton X-100 for 20 min, followed by incubation in 5 % w/v bovine serum albumin solution (BSA; Sigma Aldrich) for 1 h at room temperature with the gently shaking. To visualize the cellular morphology on the nanofiber and structural integrity, the cytoskeleton was labeled using a rabbit monoclonal Alexa Fluor® 488-conjugated anti-ß-actin antibody ([SP124], Abcam) at a 1:200 ratio in 5% w/v BSA/PBS solution, which was used to label the cytoskeleton protein overnight at 4 °C. After twice washing with PBS, Next, the samples were stained with DAPI (1:2000, Invitrogen) diluted in PBS for 10 min at room temperature at RT in the dark, after washing twice with PBS. The protein expression and signal localization were investigated using the inverted fluorescence microscope.

### Transwell migration assay

The migration of cells was assessed using Transwell chambers with 8.0 μm pore polycarbonate membrane insert (Corning, Bedford, MA, USA). The membranes were placed on the lower chamber filled with culture medium contained 10 % FBS. HUVECs (1 × 10^4^ cells/1.9 cm^2^) suspended in 200 µl serum-free culture medium were then seeded into each upper well. After 1 day of cultivation, we removed the upper well and washed the cells with PBS three times before fixed with 4% formaldehyde for 20 min at room temperature. After the fixation process, we washed the samples with PBS for two times, following gently stained with the 0.1% crystal violet for 20 min on the shaker, and then washed twice to remove the residual dye. The cell number was counted under microscopic observation.

### STZ-induced type-1 diabetic model

Healthy male adult Sprague-Dawley rats weighing 300 ± 10 g (6–8 weeks of age) were supplied by BioLASCO Taiwan Co., Ltd. (Taipei, Taiwan). All the rats were housed in an environment with a controlled temperature of 24 °C, relative humidity of 50 ± 10 %, and a fixed light/dark schedule (12 h light/12 h dark) and were allowed free access to food and water. After 3 days of adaptation, all rats were fasted for 12 h before diabetes induction. The Type 1 diabetes model was induced by a single intraperitoneal injection of 65 mg/kg STZ (streptozocin) (Sigma-Aldrich, USA) dissolved in 0.1 M citrate buffer (dissolve 0.9606 g of citric acid in 50 ml ddH2O, use 0.1 M NaOH to adjust the pH value to pH 4.5). As a control, normal rats were administered the same amount of citrate buffer alone. After 72 h of fasting, rats with a fasting blood glucose level of 200 mg/dL were considered diabetic.

### Bliss Independence Model

To mathematically distinguish between additive and synergistic effects of the combined secretome therapy *in vivo*, the expected combinatorial effect was calculated using the Bliss Independence Model. The formulation of Bliss Independence is based on monotonically increasing responses for increasing doses:

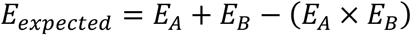

where *E_A_* and *E_B_* represent the fractional healing effects of the single treatments relative to the unhealed proportion of the control group ^[14]^. A synergistic interaction was defined when the observed therapeutic effect significantly exceeded the theoretical *E_expected_* value.

Here, we measure the effect of wound closure efficiency on day 21 in *in vivo* model. Sample groups are normalized to the control group to calculate the fractional effect as equation:

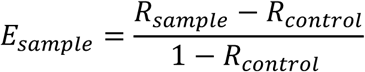

where *R_sample_* and *R_control_* represent the wound closure rate (%) of sample and negative control (bare NF scaffold) ^[15]^.

### Histopathological studies

To evaluate morphological changes, tissue regeneration, and wound-healing processes, skin specimens were prepared using distinct fixation protocols. For hematoxylin and eosin (H&E) staining, specimens were fixed in 4% formaldehyde, while those designated for Masson’s trichrome staining were fixed in Bouin’s solution. Following fixation, all tissues were embedded in paraffin and sectioned at a thickness of 3 µm. H&E staining was performed to visualize cell nuclei (blue/purple) and counterstain cytoplasmic and extracellular matrix structures (pink/red). To evaluate collagen formation and tissue fibrosis, Masson’s trichrome staining was conducted using a commercial kit provided by Rapid Science Co., Ltd. (Taichung, Taiwan) to stain collagenous connective tissue blue. All stained sections were reviewed and imaged under a light microscope. For the quantification of tissue fibrosis, the Masson’s trichrome-stained sections were analyzed using ImageJ software. The extent of collagen deposition was determined via the Color Deconvolution plugin, where the blue-stained collagen area was digitally isolated and expressed as a percentage of the total tissue area. To normalize the data across samples, the calculated area of collagen deposition was divided by the total area of black-stained nuclei.

### Immunohistochemical (IHC)

Immunohistochemical (IHC) analysis of ⍺-SMA, CD68, iNOS, COX-2, and CD206 was performed using the formalin-fixed paraffin-embedded (FFPE) tissue sections. Following deparaffinization, heat-induced antigen retrieval was conducted in a citrate buffer (pH = 6) at 95 °C for 15 min. Subsequent protein detection was carried out using the Novolink™ Polymer Detection System (Leica Biosystems, RE7140) following the manufacturer’s protocol. The specific antigenic sites were visualized via 3,3’-diaminobenzidine (DAB) chromogen, and sections were counterstained with hematoxylin.

### Data analysis

All experiments were performed in biological triplicate (n ≥ 3), and quantitative data are expressed as the mean ± standard deviation (SD). Statistical analyses were performed to evaluate significant differences among experimental groups. For datasets derived from *in vitro* experiments, a one-way ANOVA was conducted and followed by Tukey’s post hoc test for multiple comparisons. For animal experiments evaluating changes across different weeks, a two-way ANOVA was utilized to determine the main effects of the experimental groups and time points, as well as their interactions, followed by Tukey’s post hoc test for multiple comparisons. The *p*-value of less than 0.05 (*p* < 0.05) was considered statistically significant. All statistical evaluations and graphical representations were generated using GraphPad Prism 10 (GraphPad Software, San Diego, CA, USA).

## Results

### Optimization of Chinese Herbal Compounds for WJMSC Induction

To determine the optimal bioactive agents for scaffold integration, we first evaluated the cytotoxicity and functional modulation of six candidate Chinese herbal compounds including AS-IV, BA, CA, DP, FMN, and SAL. The dose-dependent toxicity of each compound was assessed in WJMSCs following a 48-hour incubation period. As demonstrated in Figure 1A, all six candidates exhibited excellent biocompatibility at concentrations up to 1 μg/mL, maintaining cell viability comparable to the control group. However, distinct toxicity profiles emerged at higher concentrations. While catechin and salidroside displayed broad safety profiles with no significant cytotoxicity up to 100 μg/mL, other compounds induced varying degrees of cell death. AS-IV maintained high viability at 10 μg/mL but exhibited a statistically significant reduction to approximately 73% at 50 μg/mL. DP and BA displayed more severe cytotoxicity which viability dropped to 9% and 21% at 50 μg/mL. In addition, DP also revealed significant decrease in cell viability with 10 μg/mL treatment. FMN revealed moderate cytotoxicity, reducing viability to less than 50% at concentrations of 50 μg/mL. Consequently, a concentration range of 0.01–1 μg/mL was established as the safe therapeutic window for subsequent functional screening.

**Figure 1.**
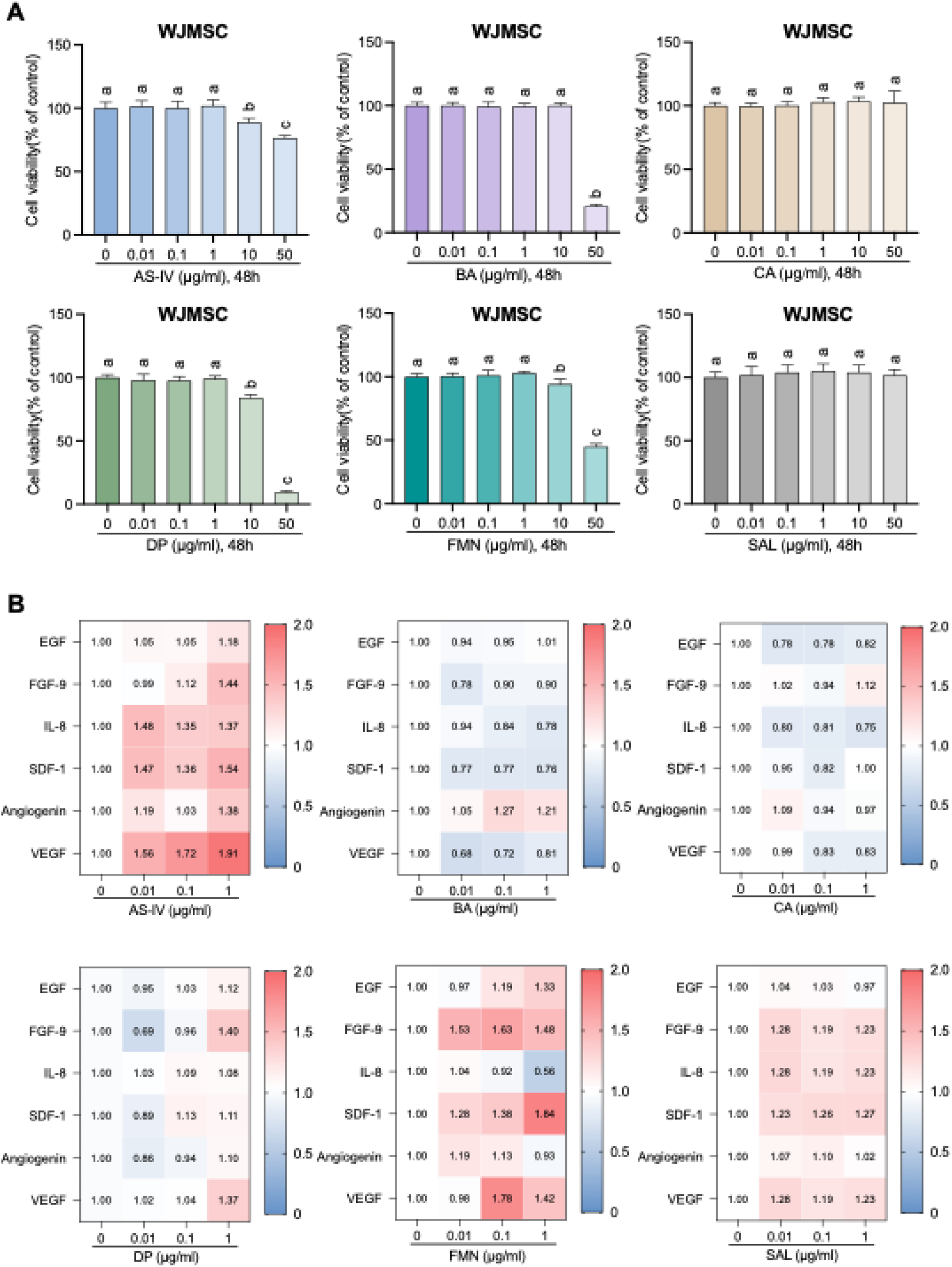
Cytokine secretion with Chinese herbal stimulation. (A) Cell viability of Wharton’s Jelly-derived Mesenchymal Stem Cells (WJMSCs) cultured with varying concentrations (0.01–50 μg/mL) of six chinese herbal compounds for 48 h, assessed via MTT assay. Compounds include astragaloside IV (AS-IV), boswellic acid (BA), catechin (CA), dracorhodin perchlorate (DP), formononetin (FMN), and salidroside (SAL). (B) Heat map quantifying the relative secretion levels of key regenerative cytokines (EGF, FGF-9, IL-8, SDF-1, Angiogenin, VEGF) from WJMSCs treated with safe concentrations (0.01–1 μg/mL) of the herbal compounds, normalized to the untreated control (0 μg/mL). Data are presented as mean ± S.D. (n = 3). \**p* < 0.05, **** *p* < 0.0001.

Following the biocompatibility assessment, the expression profiles of key wound-healing biomarkers were analyzed to identify potential inducers for the conditioned medium. The secretion levels of epidermal growth factor (EGF), fibroblast growth factor-9 (FGF-9), interleukin-8 (IL-8), stromal cell-derived factor-1 (SDF-1), angiogenin, and vascular endothelial growth factor (VEGF) were quantified. The relative expression levels were summarized in a heat map (Figure 1B), with the corresponding raw cytokine antibody array data provided in Supplementary Figure S1. The screening revealed distinct advantages among three key candidates. AS-IV elicited the most robust and broad-spectrum upregulation of regenerative factors, significantly potentiating the expression of VEGF (1.91-fold), SDF-1 (1.54-fold), FGF-9 (1.44-fold), and IL-8 (1.37-fold) at 1 μg/mL relative to controls. Formononetin demonstrated strong specific inductive potential, particularly for SDF-1 (1.84-fold) and FGF-9 (1.48-fold), effectively complementing the profile of AS-IV. Salidroside exhibited consistent upregulation across multiple markers, including VEGF (1.23-fold) and FGF-9 (1.23-fold), while maintaining the highest safety profile among all tested compounds. Conversely, BA and CA resulted in negligible modulation or downregulation of these critical markers. Based on these combined findings of safety and bioactivity, AS-IV, FMN, and SAL were selected as the primary candidates for further evaluation.

### Reversal of High-Glucose Induced Cellular Dysfunction via the Akt/eNOS Signaling

To mimic the pathological microenvironment of a diabetic wound, HDFa cells were exposed to high-glucose stress (25 mM), and the rescue capacity of the herbal-induced conditioned medium (CM) was evaluated. HDFa exposured to high-glucose conditions significantly compromised cell viability compared to the normal glucose control (Figure 2A). However, treatment with WJMSCs-CM induced by the selected herbal compounds effectively attenuated this cytotoxic effect in a concentration-dependent manner. Notably, the optimal rescue concentration varied among the candidates. The CM induced by AS-IV elicited the most potent protective effect at a low concentration of 0.01 μg/mL, restoring cell viability to near-control levels. In contrast, FMN and SAL exhibited a different dose-response profile, requiring a higher concentration of 1 μg/mL to achieve maximum cyto-protection.

**Figure 2.**
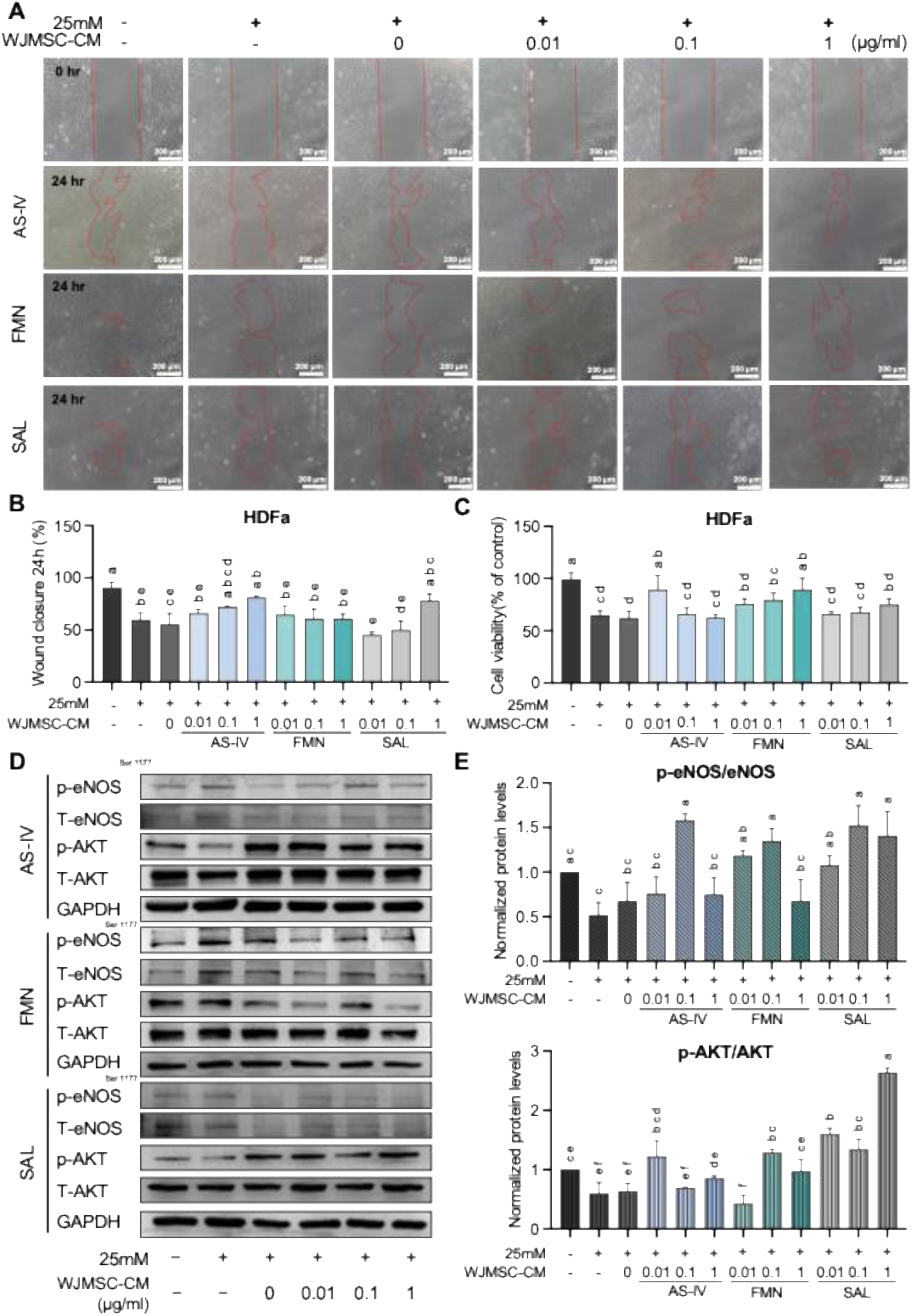
Reversal of high-glucose induced cellular dysfunction via Akt/eNOS signaling. (A) Viability of human dermal fibroblasts (HDFa) exposed to high-glucose stress (25 mM) and subsequently rescued by WJMSC conditioned medium (CM) induced by AS-IV, FMN, or SAL. (B) Representative microscopic images and corresponding quantification of HDFa migration in a scratch wound assay under high-glucose conditions with herbal-induced CM treatment at 0 h and 24 h. (C) Representative Western blot images showing the expression levels of p-eNOS, total eNOS, p-AKT, total AKT, and GAPDH (internal control) in HDFa cells following treatment. (D) Quantitative densitometric analysis of p-eNOS/eNOS and p-AKT/AKT ratios normalized to controls. Data are mean ± S.D. (n = 3). \**p* < 0.05, \*\**p* < 0.01, \*\*\**p* < 0.001, \*\*\*\**p* < 0.0001. Scale bar: 100 μm.

To further assess the functional recovery of the cells, their migratory capacity was evaluated using a scratch wound healing assay (Figure 2B). Consistent with the viability data, the hyperglycemic environment severely inhibited cell migration in the untreated control group. The cells treated with AS-IV-induced CM and FMN-induced CM were effectively accelerated the wound closure after 24 h with concentration from 0.01 μg/mL. The cells treated with SAL-induced CM didn’t show significant effect with 0.01 and 0.1 μg/mL treatment but revealed 80% wound closure at 1 μg/mL.

To elucidate the molecular mechanism underlying this cellular rescue, the activation of the PI3K/Akt/eNOS signaling, a critical pathway for cell survival and angiogenesis, was investigated via Western blotting (Figure 2C). High-glucose stress resulted in a marked suppression of the phosphorylation levels of Akt (p-Akt) and endothelial nitric oxide synthase (p-eNOS). Quantitative analysis demonstrated in Figure 2D revealed that the herbal-induced CM successfully reversed this suppression. Specifically, AS-IV-induced CM at 0.01 μg/mL significantly upregulated p-eNOS and p-Akt levels, suggesting a restoration of angiogenic signaling. FMN and SAL also induced the phosphorylation of these key proteins, with peak activation observed at 1 μg/mL. These findings indicate that the herbal-induced secretome protects cells from diabetic stress and restores functional behavior primarily by reactivating the Akt/eNOS signaling cascade.

### Enhancement of Endothelial Angiogenic Potential by Herbal-Induced Conditioned Medium

Since the typical characteristic of chronic wounds are impaired vascularization, the ability of the herbal-induced secretome to promote angiogenesis under hyperglycemic stress was evaluated using Human Umbilical Vein Endothelial Cells (HUVECs). The cyto-protective effects of the conditioned medium on endothelial cells were assessed by subjecting the cells to high glucose (25 mM). High-glucose culture conditions significantly reduced HUVECs viability compared to the normal glucose control (Figure 3A). Treatment with the herbal-induced CM reversed this impairment, though the optimal concentration varied by compound. AS-IV-induced CM maximally restored cell viability at 0.1 μg/mL, whereas FMN-induced CM exhibited a linear dose-dependent efficacy, peaking at 1 μg/mL. SAL-induced CM demonstrated peak efficacy at 0.1 μg/mL before declining at higher concentrations.

**Figure 3.**
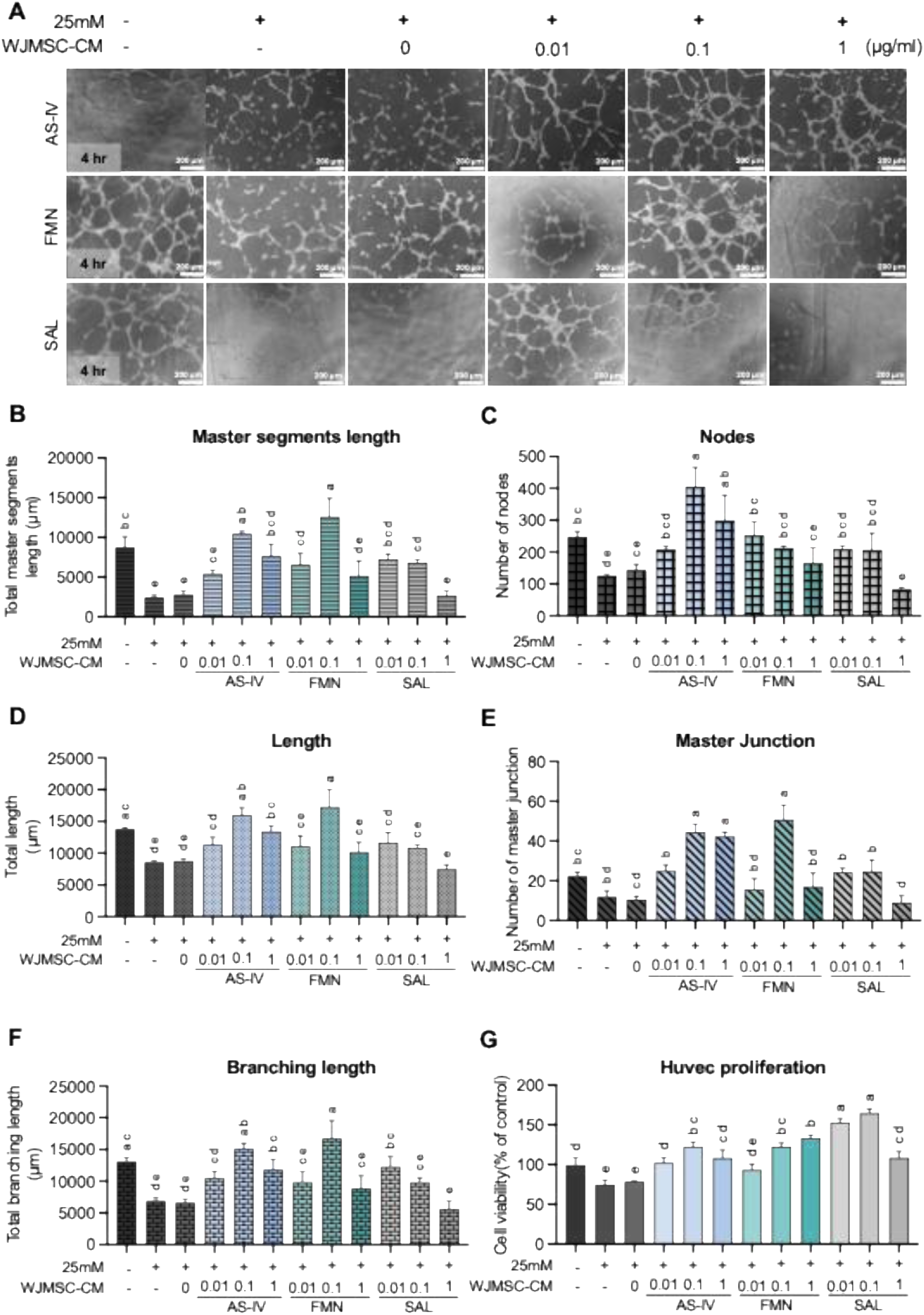
Enhancement of endothelial angiogenic potential under high-glucose stress. (A) Viability of Human Umbilical Vein Endothelial Cells (HUVECs) cultured under high-glucose conditions (25 mM) and treated with herbal-induced CM (AS-IV, FMN, SAL) at varying concentrations. (B-D), Representative images of *in vitro* tube formation by HUVECs on Matrigel after exposure to high glucose and treatment with CM induced by AS-IV (B), FMN (C), or SAL (D). Corresponding quantitative analyses of total branching length, number of master junctions, number of nodes, and total master segments length are provided below the images. Data are mean ± S.D. (n = 3). \**p* < 0.05, \*\**p* < 0.01, \*\*\**p* < 0.001, \*\*\*\**p* < 0.0001. Scale bar: 100 μm.

To evaluate functional angiogenesis, an *in vitro* tube formation assay was performed. Under high-glucose conditions, HUVECs failed to form organized capillary-like structures, exhibiting fragmented networks with significantly reduced branching points and master junctions. The application of herbal-induced CM effectively rescued this phenotype, promoting the formation of intact, complex meshes. Quantitative analysis of the topological parameters revealed distinct potency profiles. For AS-IV, the 0.1 μg/mL induction concentration yielded the most robust angiogenic network, significantly increasing the number of master junctions, nodes, and total branching length to levels comparable to the negative control (Figure 3B). FMN also displayed strong angiogenic potential, with the 0.1 μg/mL concentration inducing the highest density of vascular structures (Figure 3C). Interestingly, while the 1 μg/mL concentration of FMN maximized cell viability, it was less effective at promoting tube formation compared to the 0.1 μg/mL dose, suggesting a specific window for morphogenic signaling. SAL exhibited a different profile, with peak tube formation observed at the lowest concentration of 0.01 μg/mL (Figure 3D). Higher concentrations of SAL resulted in a regression of the vascular network, indicating a narrower therapeutic index for angiogenic stimulation compared to AS-IV and FMN. Collectively, these data demonstrate that the herbal-induced WJMSCs-CM can effectively overcome the anti-angiogenic effects of a high-glucose microenvironment, restoring the capacity of endothelial cells to organize into functional vascular networks.

### Nanofiber-based scaffold fabrication and characterization

Core-shell nanofibrous scaffolds were fabricated using a polyvinyl alcohol (PVA) and gelatin polymer system via coaxial electrospinning (Figure 4A). To achieve uniform nanofibers, the polymer flow rate was systematically varied. Result from scanning electron microscopy (SEM) showed that the flow rate directly influenced fiber morphology (Figure 4B). Higher flow rates (0.3–0.4 mL/h) produced thicker, merged fibers with peak diameters between 500 and 600 nm. Decreasing the flow rate to 0.06 mL/hr produced finer, smoother fibers, shifting the peak diameter to approximately 250 nm (Figure 4C). This diameter reduction corresponded to a decrease in overall scaffold porosity, dropping from 85% at 0.4 mL/h to 71% at 0.06 mL/h (Figure 4D). To balance the fiber diameter with adequate porosity for cellular infiltration, an intermediate flow rate 0.06 mL/hr was selected for all subsequent scaffold fabrications.

**Figure 4.**
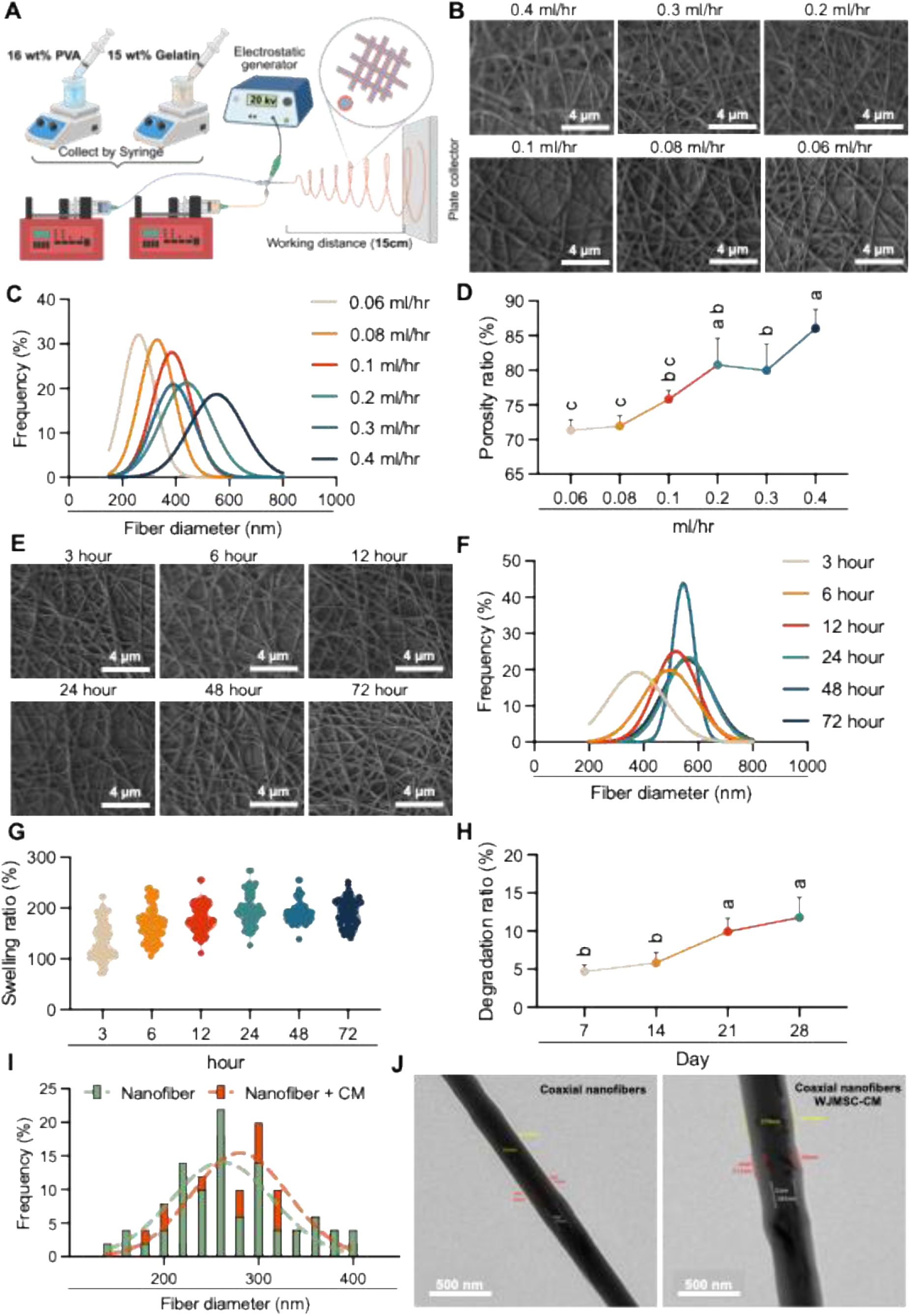
Fabrication and characterization of coaxial electrospun nanofibrous scaffolds. (A) Schematic illustration of the core-shell nanofiber fabrication process via coaxial electrospinning using PVA (core) and gelatin (shell). (B) Representative scanning electron microscopy (SEM) images of nanofibers fabricated at varying flow rates (0.06–0.4 mL/h). (C) Fiber diameter distribution curves corresponding to the varying flow rates. (D) Quantitative analysis of scaffold surface porosity across different flow rates. (E) Representative SEM images of the optimized nanofiber scaffolds undergoing hydration and swelling over 72 h. (F) Fiber diameter distribution curves tracking swelling behavior over time. (G) Quantification of the temporal swelling ratio of the scaffolds. (H) *In vitro* degradation profile of the nanofiber scaffolds in PBS over 28 days. (I) Diameter distribution of unloaded nanofibers versus nanofibers loaded with CM. (J) Transmission electron microscopy (TEM) images confirming the core-shell biphasic structure of the unloaded (left) and CM-loaded (right) nanofibers. Data are mean ± S.D. (n = 3). ns: no significant, \**p* < 0.05, \*\**p* < 0.01, \*\*\**p* < 0.001, \*\*\*\**p* < 0.0001. (Figure 4A created in BioRender. Huang, L. C. (2026))

To evaluate the structural stability of the scaffolds in an aqueous environment, we measured the fiber swelling ratio and degradation behavior via nanofibrous diameter changes (Figure 4E-G). The nanofibrous showed progressive swelling over 24 h, and maintain the swelling ratio at around 120% (Figure 4G). In addition, we further evaluated the degradation ratio, and verified a low mass loss after 28 days incubation, with the scaffolds only revealed ∼12% shrinkage by Day 28 (Figure 4H). To evaluate the core-shell structure and the encapsulation ability, nanofibrous encapsulating with and without condition medium were accessed by Transmission electron microscopy (TEM). The fibrous loading with condition medium revealed slightly thicker diameter compared to the nanofibrous without CM (Figure 4I). The biphasic structure was illustrated by a distinct core and a continuous shell (Figure 4J).

### Cellular behavior co-culture on nanofiber scaffold

The release profile of the encapsulated secretome was quantified over 14 days (Figure 5A). The core-shell scaffolds exhibited a sustained, continuous accumulation profile without a rapid initial burst for all key regenerative cytokines evaluated. By Day 14, across AS-IV, FMN, SAL induced groups, the scaffolds successfully released distinct concentrations of biologically active factors, including FGF-9 (914.72–1382.11 pg/mL), VEGF (9.91–16.81 pg/mL), IL-8 (92.87–111.22 pg/mL), EGF (9.91–16.81 pg/mL), Angiogenin (5170.27–6036.00 pg/mL), and SDF-1 (712.00–913.71 pg/mL). These robust physical and sustained release characteristics confirm that the core-shell scaffold is highly effective at providing long-term, localized delivery of the incorporated multi-protein therapeutic payload. Cellular attachment and morphological spreading on the electrospun scaffolds were evaluated using SEM and immunofluorescence staining for β-actin (Figure 5B, 5D). Cells adhered to the nanofibrous architecture and integrated into the porous network under normal conditions. Exposure to high-glucose stress visibly impaired cellular spreading and reduced cytoskeletal extensions on the unloaded scaffolds. Conversely, cultivation on scaffolds loaded with the herbal-induced CM (AS-IV, FMN, and SAL) reversed this morphological impairment, restoring robust cytoskeletal organization and widespread cellular coverage across the scaffold surface. This structural recovery demonstrates that the released CM actively protects cellular architecture against hyperglycemic stress. To quantify the cyto-protective effects of the functionalized scaffolds, the viability of human dermal fibroblasts (HDFa) and human umbilical vein endothelial cells (HUVECs) was measured under high-glucose conditions (25 mM). High glucose exposure significantly reduced cell viability on unloaded scaffolds (Figure 5C). Specifically, HDFa viability decreased to approximately 48% of the normal control, and HUVECs viability decreased to roughly 80%. Culturing the cells on scaffolds loaded with herbal-induced CM significantly restored viability in both cell lines. For HDFa, viability increased to 122%, 141%, and 139% on the A, F, and S-loaded scaffolds. Similarly, HUVECs viability was restored to 99%, 119%, and 149% across the respective CM groups. These data indicate that the sustained delivery of the primed secretome from the core-shell scaffold successfully preserves both fibroblast and endothelial cell viability in a diabetic-like microenvironment.

**Figure 5.**
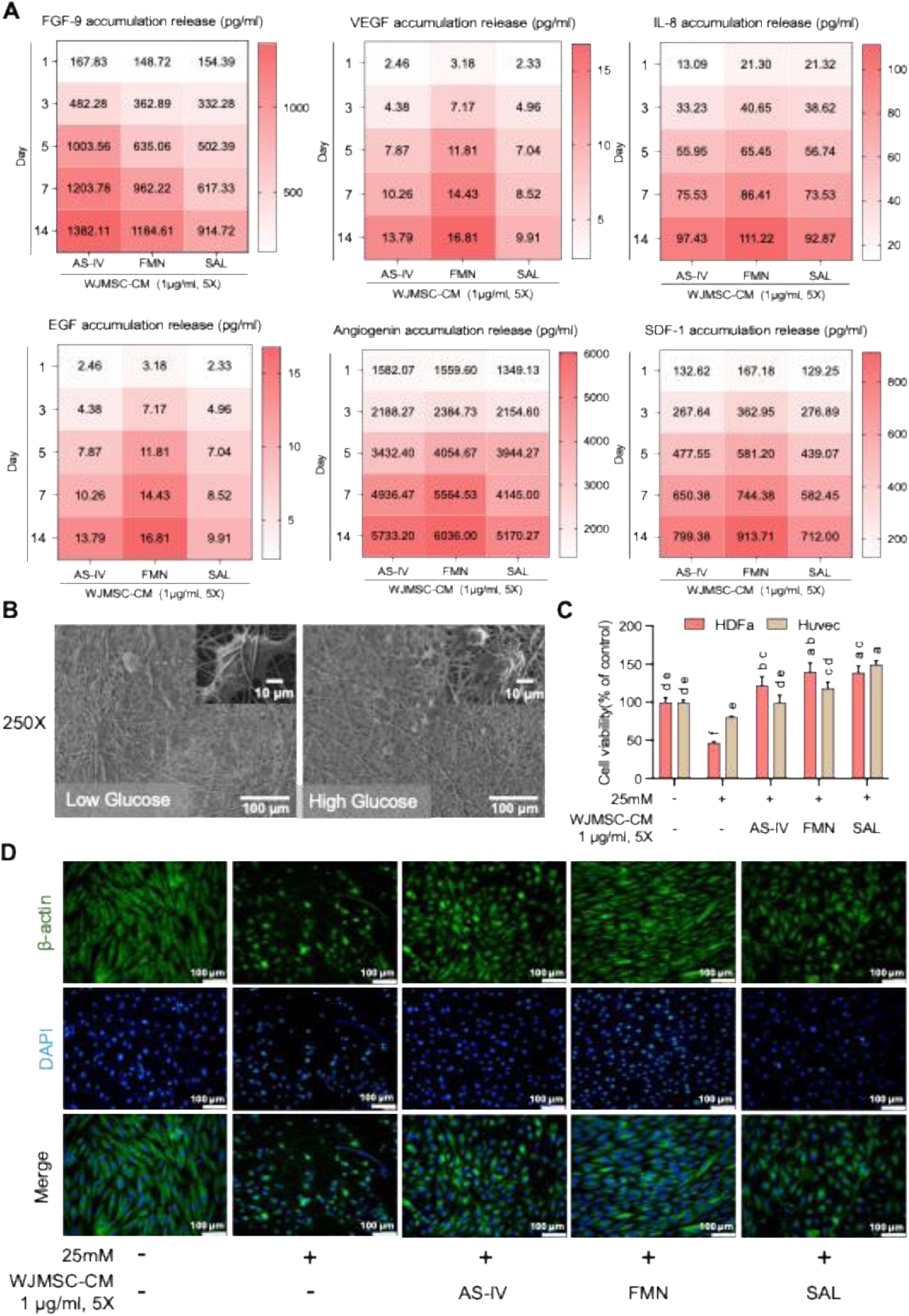
Sustained release kinetics and cellular behavior on CM-loaded nanofibrous scaffolds. (A) Cumulative *in vitro* release profiles of key regenerative cytokines (FGF-9, VEGF, IL-8, EGF, Angiogenin, SDF-1) from the core-shell scaffolds loaded with AS-IV, FMN, and SAL-induced CM over 14 days. (B)Representative SEM images demonstrating cellular attachment and morphological spreading on unloaded (left) and CM-loaded (right) nanofiber scaffolds. (C) Viability of HDFa and HUVECs co-cultured on the functionalized scaffolds under high-glucose stress (25 mM). (D) Immunofluorescence staining of cells on the scaffolds evaluating cytoskeletal organization (β-actin, green) and nuclei (DAPI, blue) under high-glucose conditions with or without localized CM delivery. Different letters indicate statistically significant differences at *p* < 0.05.

### Functional Matrix Secretion and Endothelial Migration on Scaffolds

To evaluate the migratory capacity of endothelial cells in a high-glucose microenvironment (25 mM), Transwell migration assays were performed using HUVECs seeded on the core-shell scaffolds (Figure 6A). Quantitative analysis of HUVECs migration demonstrated that high glucose severely impaired cell motility on the scaffold with and without CM. Local delivery of the herbal-primed secretome significantly rescued this migratory dysfunction. While both the AS-IV and SAL-primed secretomes moderately promoted migration compared to bare scaffolds, the FMN-loaded scaffold group elicited the most pronounced rescue effect, driving relative cell migration significantly beyond that of the high-glucose control group. This enhanced migratory behavior confirms that the sustained release of FMN-induced factors provides highly active chemotactic signals for endothelial cells under hyperglycemic stress. We further accessed the tissue remodeling marker matrix metalloproteinase-1 (MMP-1) and pro-collagen Iα secretion by HDFa cells on the scaffold under high-glucose conditions (Figure 6B). High-glucose stress significantly suppressed the synthesis of pro-collagen I α1 on bare scaffolds compared to normal controls. Co-culture on scaffolds functionalized with the herbal-induced CM rescued this secretory deficit, with the AS-IV and FMN groups restoring pro-collagen levels to near-normal physiological baselines. Concurrent analysis of matrix metalloproteinase-1 (MMP-1), an enzyme essential for early wound debridement and matrix turnover, revealed a significant upregulation in secretion for all CM-loaded scaffolds, peaking in the AS-IV and SAL group.

**Figure 6.**
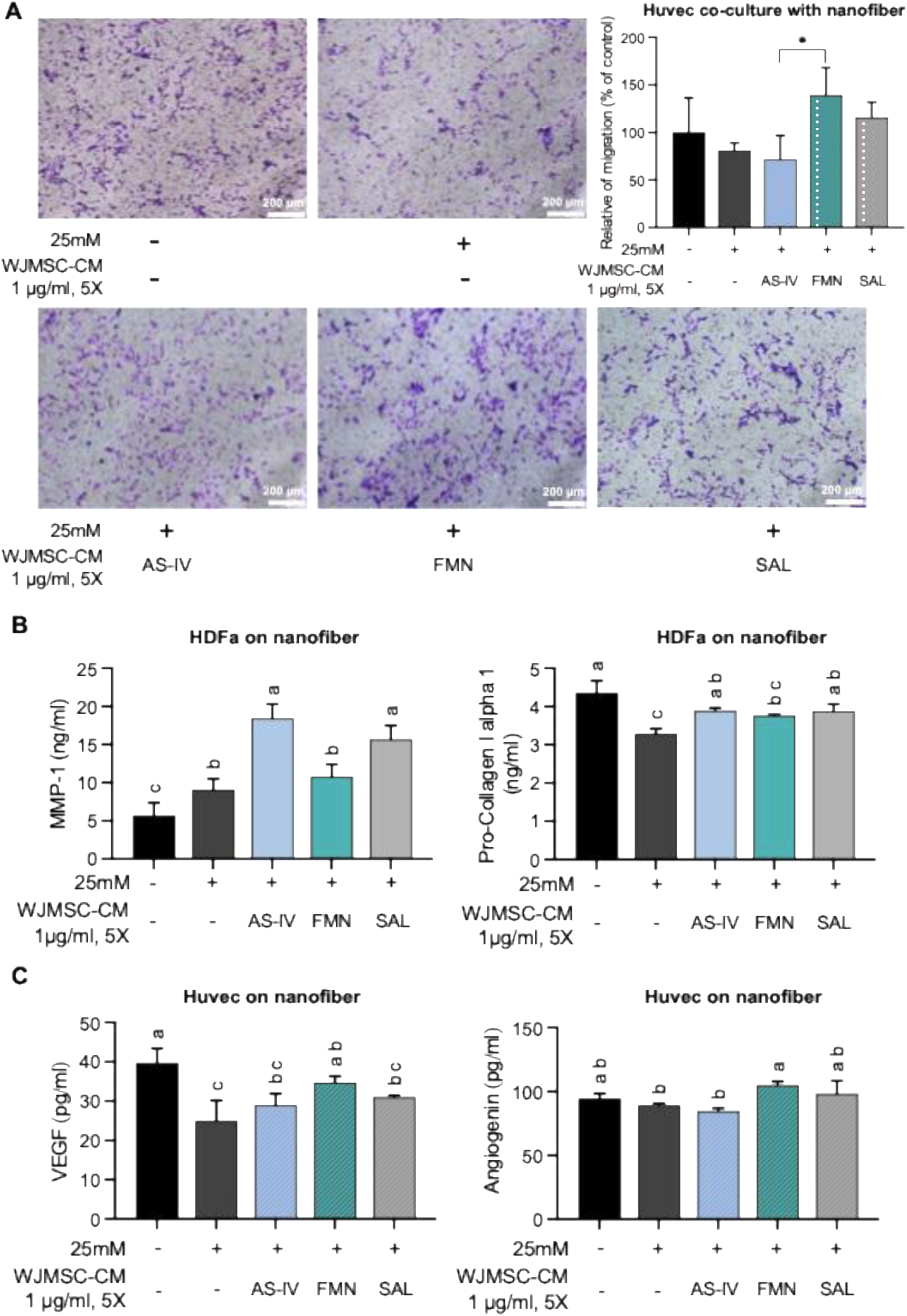
Functional matrix secretion and endothelial migration on functionalized scaffolds. (A) Representative images of HUVECs Transwell migration assays under high-glucose stress across different scaffold groups. (B) Quantification of of matrix metalloproteinase-1 (MMP-1) and pro-collagen Iα secretion by HDFa cells cultured on bare or CM-loaded scaffolds under high-glucose stress. (C) Quantification of angiogenin and VEGF secretion by HUVECs co-cultured on the scaffolds under high-glucose stress. Data are presented as mean ± S.D. (n = 3). Different letters indicate statistically significant differences (*p* < 0.05) determined by one-way ANOVA followed by Tukey’s post hoc test.

To determine if the scaffolds could sustain angiogenic signaling, the synthesis of potent vascular growth factors by co-cultured HUVECs under high-glucose stress was quantified (Figure 6C). Exposure to high glucose markedly reduced the baseline secretion of both angiogenin and vascular endothelial growth factor (VEGF) on the scaffold without CM. Endothelial cells cultured on FMN-induced CM-loaded scaffolds displayed the most robust upregulation of angiogenin and VEGF. The SAL-induced CM also supported high levels of factor synthesis, whereas the AS-IV-induced CM produced a more moderate angiogenic response. These findings confirm that the functionalized scaffolds actively promote both fibroblast-mediated ECM remodeling and endothelial cell migration required for neo-vascularization under high-glucose condition.

### Wound healing efficiency on type 1-diabetic rat model

To validate the establishment of a chronic diabetic microenvironment, we confirmed the induction of Type-1 diabetes via a single intraperitoneal injection of STZ (Figure 7A). Successful model establishment was defined by fasting blood glucose levels exceeding 200 mg/dL at 72 hours post-injection, which remained sustained throughout the 21-day study period (Figure 7B). Full-thickness cutaneous wounds (10 × 10 mm) were created on the dorsum of the diabetic rats to evaluate the therapeutic efficacy of the CM-loaded coaxial scaffolds. Therapeutic efficacy was then evaluated using a full-thickness excision wound model. We monitored the wound closure continuously (Figure 7C) and quantitatively analyzed over 21 days (Figure 7D). On Day 7, different wound-healing kinetics were observed among the single-treatment groups; the NF-AS-IV group promoted rapid initial contraction (60.2 ± 6.8 %), whereas the NF-FMN group exhibited a delayed early response (35.1 ± 5.2 %). By Day 14, healing trajectories across all scaffold-treated groups began to converge, with the combination NF-AF group exhibiting the highest average closure (85.6 ± 4.9 %). By Day 21, the therapeutic benefit of co-delivery became highly pronounced. The combination NF-AF group achieved near-complete wound closure (98.2 ± 1.5 %), which was statistically superior to both the individual NF-AS-IV (89.2 ± 5.5 %) and NF-FMN (90.1 ± 4.8 %) treatment groups (*p* < 0.05). In comparison, wounds treated with the bare NF scaffold (85.4 ± 5.1 %) or standard controls plateaued with incomplete closure. These outcomes demonstrate that while the individual primed secretomes promote distinct phases of repair, their sustained co-delivery from the coaxial scaffold is required to trigger the synergistic cascade necessary for complete diabetic wound closure. Crucially, mathematical evaluation using the Bliss Independence Model confirmed a synergistic rather than merely additive interaction. While the theoretical additive wound closure for the combined treatment was calculated to be 92.67%, the observed closure in the NF-AF group reached 98.2%, definitively establishing the synergistic potency of co-delivering AS-IV and FMN-primed secretomes.

**Figure 7.**
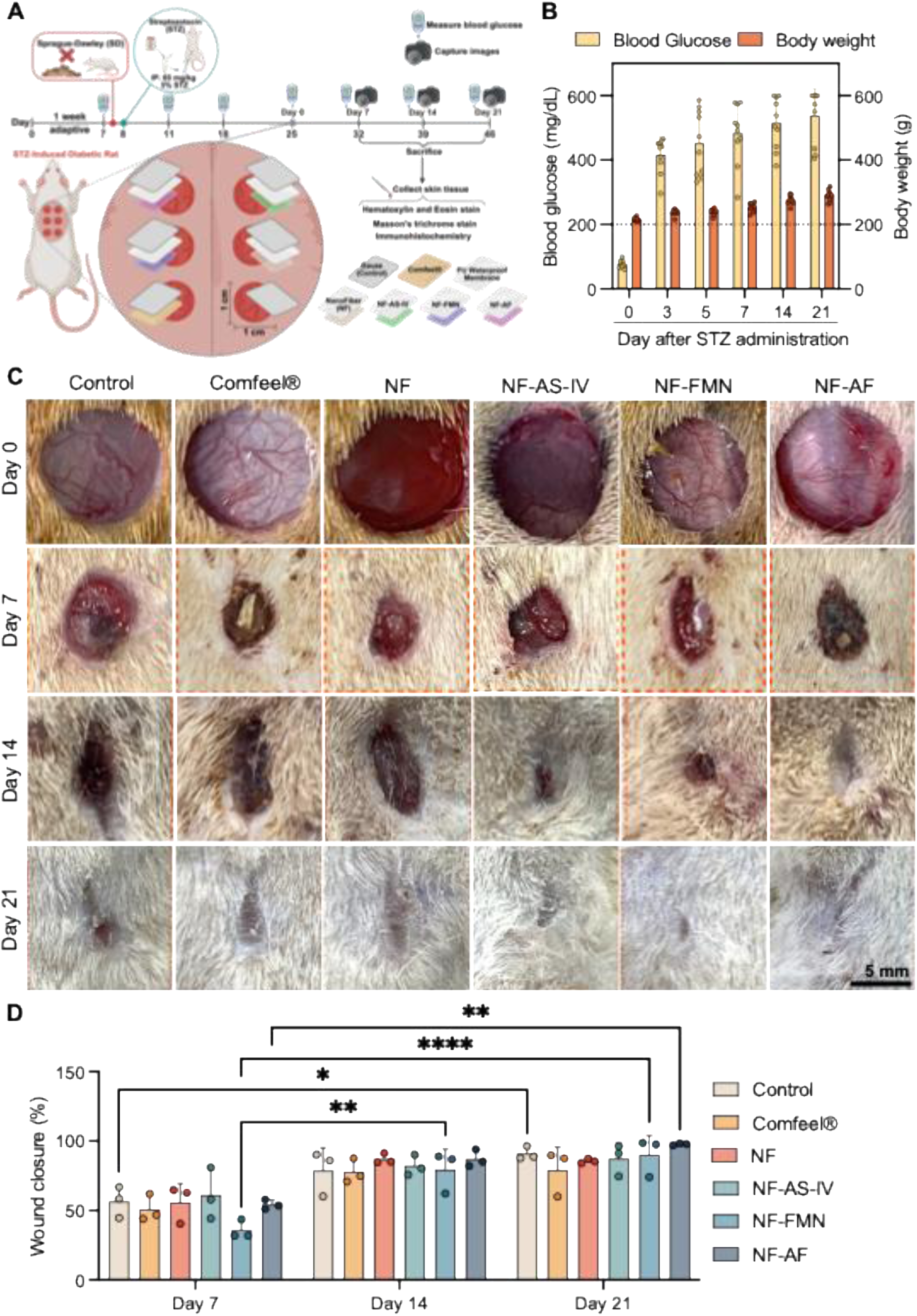
*In vivo* wound healing efficacy of functionalized scaffolds in diabetic rat model. (A) Schematic overview of the experimental timeline design of STZ-induced Type 1 diabetic rat model generation, and treatment evaluation (B) Monitoring of blood glucose levels and body weight throughout the 21-day study period, confirming sustained hyperglycemia. (C) Representative macroscopic photographs of the full-thickness cutaneous wounds receiving different treatments (Control, Comfeel®, NF, NF-AS-IV, NF-FMN, NF-AF) on days 0, 7, 14, and 21 post-wounding. (D) Quantitative analysis of the temporal wound closure percentage. Data are mean ± S.D. (n = 3). Different letters indicate statistically significant differences at *p* < 0.05. Scale bar: 5 mm. (Figure 7A created in BioRender. Huang, L. C. (2026))

### Histological evaluation of epidermal and dermal tissue regeneration

To evaluate the structural regeneration of the injured tissue under diabetic conditions, histological sections of the wound beds were stained with hematoxylin and eosin (H&E) on days 7, 14, and 21 (Figure 8A). In H&E staining, deep purple coloration indicates dense nuclear material, representing inflammatory cell infiltration and early, highly cellular granulation tissue. Conversely, varying shades of pink indicate cytoplasm and the deposition of ECM proteins like collagen. On Day 7, all injured groups (Control, Comfeel, NF, NF-AS-IV, NF-FMN, and NF-AF) displayed predominantly purple-stained wound beds, indicative of heavy inflammatory infiltration. By Day 14, the untreated Control and Comfeel groups retained wide, unhealed gaps with dense purple regions. The bare NF group exhibited early-epidermal bridging but maintained a disorganized sub-structure. The single-treatment groups (NF-AS-IV and NF-FMN) began transitioning to a pink-stained dermal architecture. The combined NF-AF group demonstrated the most advanced repair, forming a continuous epidermal layer over mature ECM structure. By Day 21, while Control and bare NF wounds still exhibited irregular tissue integration, the NF-AF treated wounds achieved complete epidermal coverage and dermal structures that most closely resembled the uninjured Healthy group.

**Figure 8.**
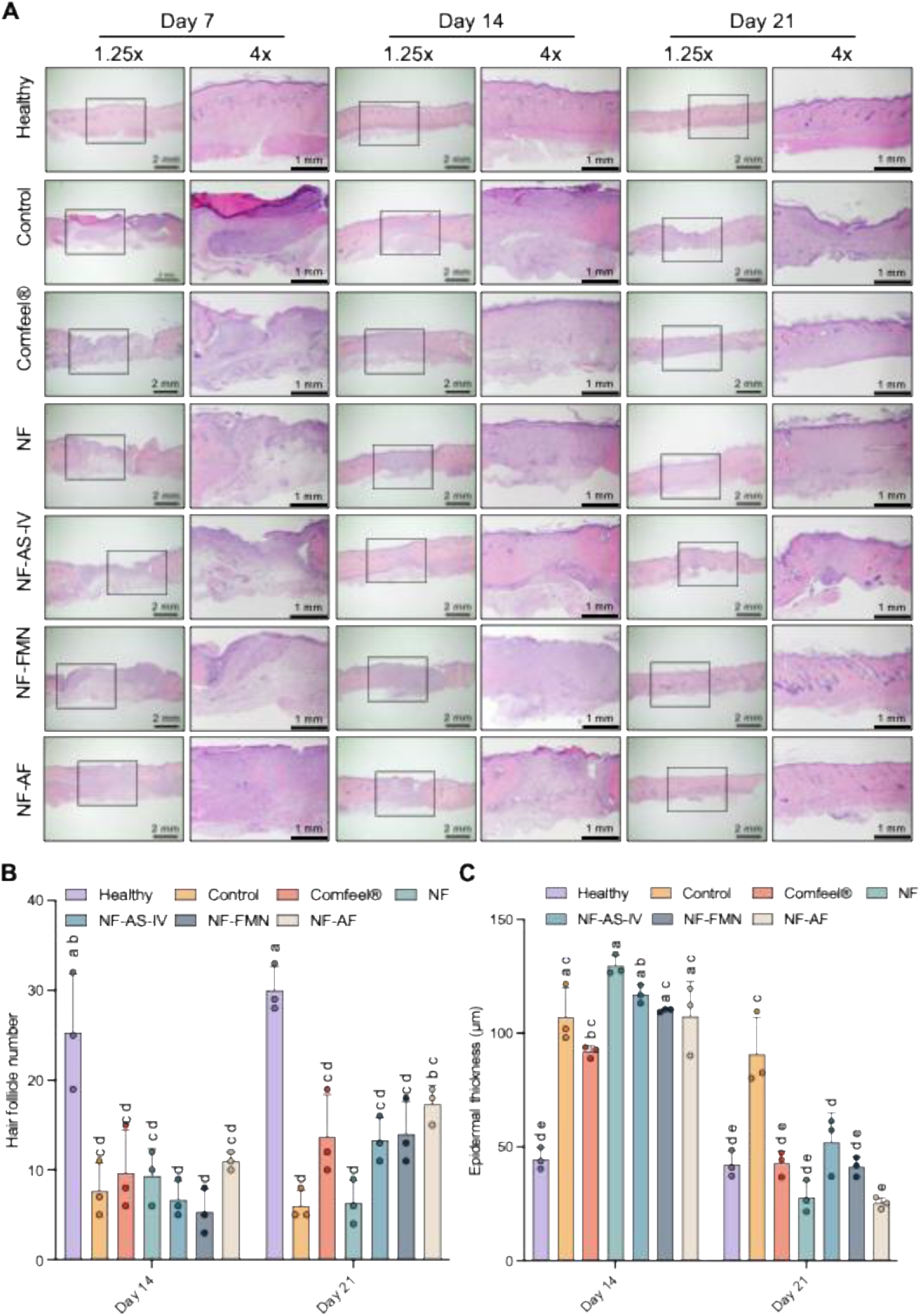
Histological assessment of epidermal and dermal tissue regeneration. (A) Representative Hematoxylin and Eosin (H&E) stained images of the regenerating wound beds from different treatment groups on days 7, 14, and 21. Black boxes denote the regions magnified in the adjacent 4x panels. (B) Quantification of the number of regenerating hair follicles within the wound bed on days 14 and 21. (C) Measurement of epidermal thickness across the treatment groups to evaluate re-epithelialization quality. Data are mean ± S.D. (n = 3). Different letters indicate statistically significant differences at *p* < 0.05.

To assess functional tissue restoration versus fibrotic scar formation, the number of newly formed hair follicles within the repaired dermis was quantified (Figure 8B). The uninjured healthy group maintained physiological follicle counts (∼30). Across days 14 and 21, the untreated control and bare NF groups exhibited minimal hair follicle neo-genesis (counts < 10 by day 21), indicating healing primarily via scar tissue. The comfeel and single secretome treatments (NF-AS-IV and NF-FMN) promoted moderate appendage recovery, reaching approximately 13–14 follicles by day 21. The combined NF-AF scaffold group achieved the highest follicle regeneration among the injured cohorts (∼17 follicles) by day 21 (*p* < 0.05, designated by distinct letters), indicating active stimulation of true dermal appendage regeneration.

To characterize tissue maturation, epidermal thickness across the healing wound beds was measured (Figure 8C). On Day 14, all injured groups exhibited pronounced pathological epidermal hypertrophy compared to the healthy baseline (∼44.7 µm), with thicknesses ranging between 90 and 130 µm due to hyper-proliferation. By day 21, healing trajectories diverged significantly. The control group maintained a persistently thick epidermis (∼90.8 µm), indicating delayed remodeling. The comfeel, NF-AS-IV, and NF-FMN groups reduced their epidermal thickness to approximately 45–50 µm. Notably, both the bare NF and the NF-AF groups demonstrated the most substantial structural remodeling, declining to approximately 30 µm and 28 µm, respectively (*p* < 0.05, distinct letters), thereby effectively matching the physiological thickness of the healthy control.

To assess the extent of extracellular matrix (ECM) synthesis and structural maturation, we evaluated the regenerating wound tissues using Masson’s trichrome staining, where mature collagen fibers are stained in blue and cellular cytoplasm/keratin are stained in red and cell nuclei in dark brawn (Figure 9A). In the uninjured healthy group, the dermis was characterized by densely packed, well-organized blue collagen structure. On day 7, the wounds from untreated control and bare NF exhibited sparse and highly disorganized collagen networks, indicating delayed matrix synthesis. The comfeel group demonstrated a rapid spike in the normalized collagen area (Figure 9B). However, histological observation revealed this early deposition as dense, amorphous fibrotic clumps rather than structured tissue. The secretome-treated groups (NF-AS-IV, NF-FMN, and NF-AF) displayed moderate early collagen deposition with emerging structural alignment. By day 14 and progressing to day 21, the trajectories of matrix remodeling diverged based on the applied treatment. The untreated control and comfeel® groups maintained irregular, heavily aggregated collagen structures typical of fibrotic scar formation. The bare NF wounds exhibited fluctuating collagen levels that lacked uniform structural integration with the surrounding healthy tissue. Conversely, the single secretome treatments (NF-AS-IV and NF-FMN) guided more aligned collagen fibrillogenesis compared to the non-biological controls. By day 21, the combined NF-AF group demonstrated the most advanced matrix maturation among all injured cohorts. While the quantitative normalized collagen area stabilized at intermediate, physiological levels (*p* < 0.05, denoted by distinct letters), the visual architecture of the blue-stained matrix in the NF-AF group transitioned into thick, uniformly distributed, and continuous wavy bundles. This structural remodeling closely mirrored the physiological ECM organization of the Healthy baseline, indicating that the co-delivery of the primed secretomes prevents bulk fibrotic scarring and instead promotes functional, high-quality dermal maturation.

**Figure 9.**
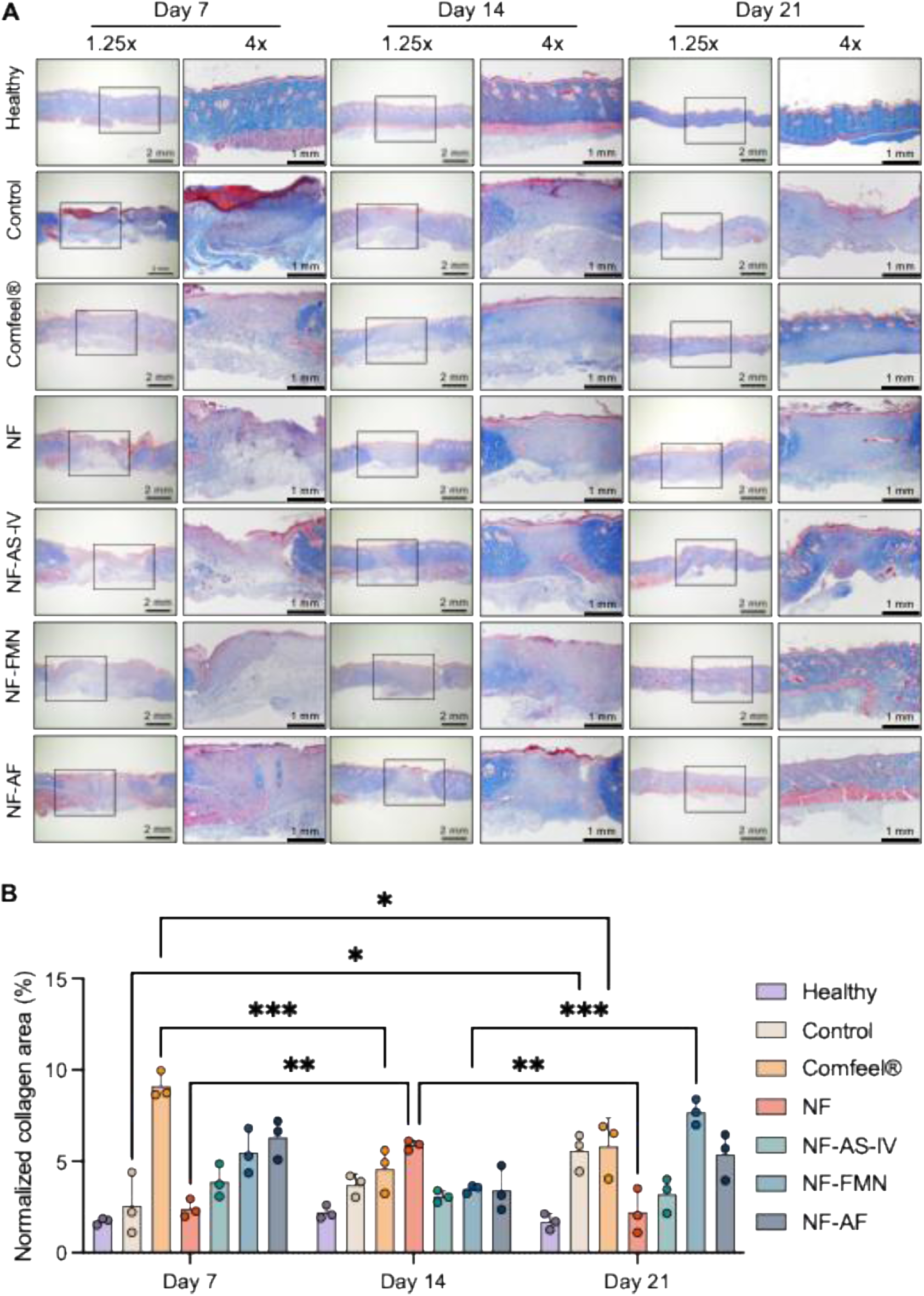
Evaluation of collagen deposition and matrix maturation *in vivo*. Representative Masson’s trichrome stained histological sections of the wound beds on days 7, 14, and 21, visualizing collagen deposition (blue). Black boxes denote the regions magnified in the adjacent 4x panels. (B) Quantitative analysis of the normalized collagen area across the different treatment groups over time. Data are mean ± S.D. (n = 3). Different letters indicate statistically significant differences at *p* < 0.05.

To evaluate the initial inflammatory state within the diabetic wound bed, IHC staining for pan-macrophage (CD68) and pro-inflammatory markers (iNOS, COX-2) was visualized on day 7 (Figure 10A). The untreated control and comfeel groups displayed dense, widespread positive staining for these markers, indicating severe inflammation. Concurrently, to visually assess early tissue regeneration and the formation of granulation tissue, α-smooth muscle actin (α-SMA) which known as the biomarker of myofibroblast formation was tracked across days 7, 14, and 21 (Figure 10B). While control wounds showed sparse early α-SMA expression, the secretome-loaded scaffolds (NF-AS-IV, NF-FMN, and NF-AF) promoted a robust, early presence of α-SMA-positive granulation tissue by day 7. Quantitative analysis of the day 7 positive staining intensity (Figure 10C) confirmed the visual observations of the inflammatory state. The secretome-loaded scaffolds significantly suppressed the expression of CD68, iNOS, and COX-2 compared to the non-biological controls (*p* < 0.05, designated by distinct letters), indicating effective dampening of the early pro-inflammatory microenvironment. Subsequent temporal quantification of α-SMA expression (Figure 10D) revealed distinct regenerative kinetics. The NF-AF group exhibited an early peak in α-SMA on day 7, indicating rapid initiation of granulation tissue formation. In contrast, the untreated control and comfeel groups displayed delayed regenerative responses, with α-SMA spiking massively later on day 14. Following their early regenerative peak, the α-SMA levels in the NF-AF treated wounds progressively resolved by day 21 to baseline levels. Finally, to confirm the sustained anti-inflammatory effect during the progression of tissue repair, the expression of CD206 was quantified on days 7 and 14 (Figure 10E). While the comfeel treatment induced an immediate, atypical spike in CD206 expression on day 7, it failed to maintain a progressive trajectory. In contrast, the secretome-loaded groups demonstrated a steady, temporally sustained upregulation of CD206 from day 7 to day 14. By day 14, the combined NF-AF group reached elevated CD206 levels that significantly outperformed both the untreated control and bare NF groups (p < 0.05, distinct letters). This progressive upregulation serves as a definitive confirmation of the sustained anti-inflammatory effect promoted by the coaxial scaffold during the critical tissue repair phase.

**Figure 10.**
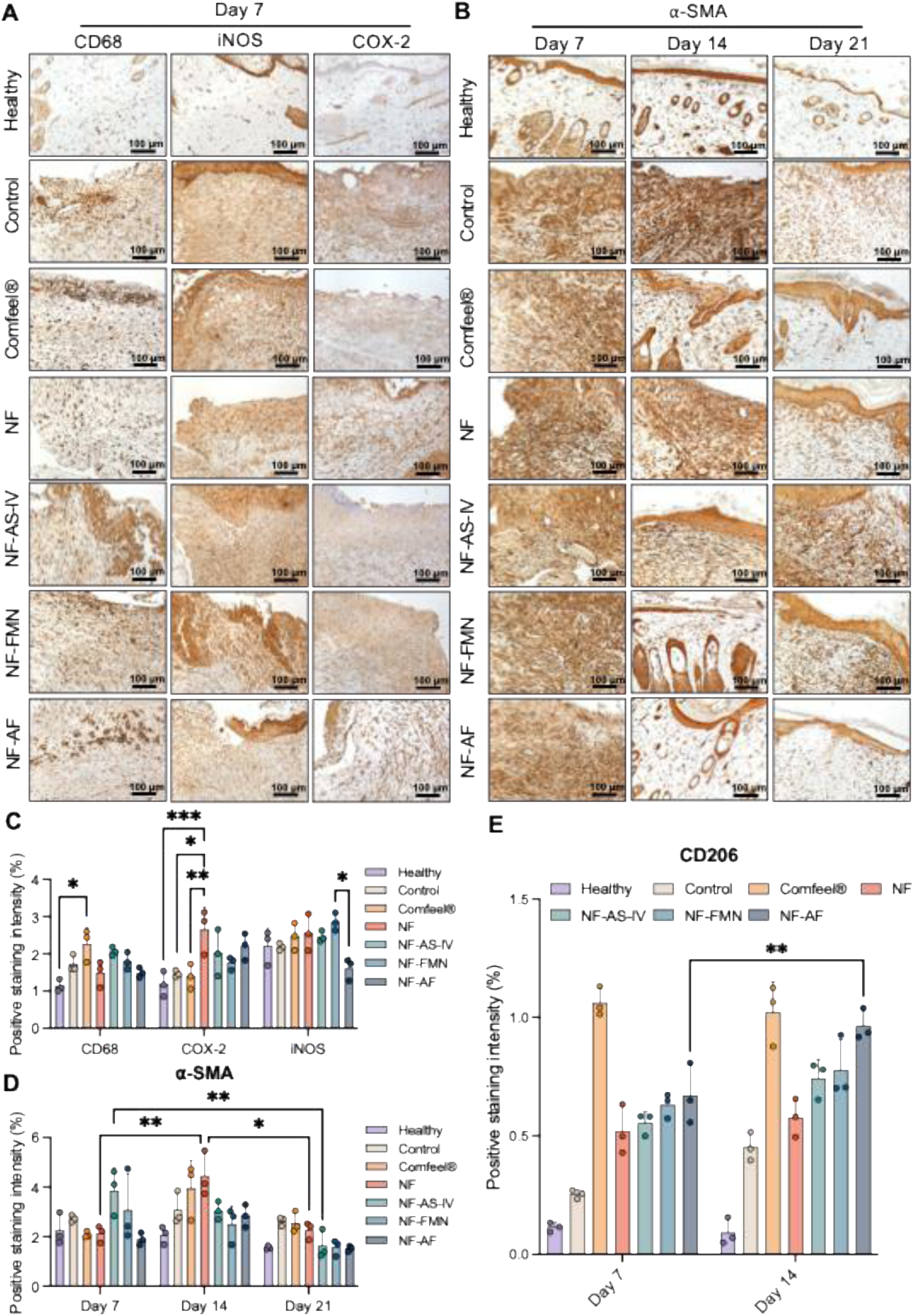
Immunomodulation and neo-vascularization promoted by CM-loaded scaffolds *in vivo*. (A) Representative immunohistochemical (IHC) staining images of the wound bed tissues on Day 7 evaluating pan-macrophage infiltration (CD68) and pro-inflammatory markers (iNOS, COX-2). (B) Representative IHC images assessing the temporal expression of the mature vascular marker α-smooth muscle actin (α-SMA) on days 7, 14, and 21. (C) Quantitative analysis of the positive staining intensity for CD68, COX-2, and iNOS on Day 7. (D) Quantitative temporal analysis of α-SMA positive staining intensity. (E) Quantitative analysis of the anti-inflammatory, pro-remodeling M2 macrophage marker CD206 on days 7 and 14. Data are mean ± S.D. (n =3). Different letters indicate statistically significant differences at *p* < 0.05.

## Discussion

Diabetic chronic wounds are among the most formidable complications of diabetes in a clinical setting ^[16]^. The hyperglycemic milieu presents barriers in the microenvironment, contributing to neuropathy and sustained chronic inflammation, which generally impede fibroblast response to severe injury, proliferation, and migration in pathological conditions, thereby preventing long-term wound healing ^[17]^. Therefore, the medical field is in immediate need of groundbreaking regenerative medicine techniques that can modify the milieu defined by ongoing inflammation and impaired recovery ^[18]^. Considering that the repair mechanism is significantly dependent on the collaborative function of many growth factors and cytokines, researchers have focused on WJMSCs, which known for their robust paracrine capabilities. These cells operate as meticulous biological manufacturing units, perpetually releasing bioactive compounds such as cytokines, growth factors, and exosomes ^[19]^. The conditioned medium from WJMSCs fortified with these elements offers an essential physicochemical mechanism for closing the biochemical signaling void in the repair of diabetic wounds.

Building upon this, we activated the cell reactions via TCM treatment with the strong paracrine potential from WJMSCs, capable of continuously secreting various pro-repair cytokines, growth factors, and exosomes ^[19]^. In this case, we elevate the efficacy of WJMSCs-CM to a new level. The preconditioning strategy in this work is a novel approach that combines TCM and stem cell biology. We screened six well-known TCM that are used to treat diabetic chronic wounds in clinical trials, and applied them to the WJMSCs. Demonstrating that concentrations up to 1 µg/ml do not compromise WJMSCs viability establishes a favorable therapeutic window for these six TCMs. Most importantly, following the intervention of TCM, the WJMSCs paracrine behaviors were significantly regulated, including bioactive components such as growth factors and cytokines. The secretory profile exhibits distinct qualitative and quantitative changes. This significant finding not only demonstrates that TCM can act as a microenvironment molecular switch for remodeling the paracrine functions of stem cells, but it also suggests that this type of optimized restriction culture medium for traditional Chinese medicine can be used to accelerate diabetic wound healing, providing the critical physicochemical mechanism basis.

After establishing the cytotoxicity of TCM and WJMSCs-CM, this study assessed the WJMSCs-CM regulating efficacy of biological behavior on HDFa. The cell viability results showed significant increased proliferation rate of HDFa in co-cultured with WJMSCs-CM compared to the negative control under high glucose conditions. This demonstrated that pre-treatment with the TCM successfully primed WJMSCs paracrine secretion, thereby supplying HDFa cells with a high active biological repair microenvironment, and thus enhancing the efficient of wound healing. In a scratch assay that mimicked a skin wound, the WJMSCs-CM promoted the HDFa cells migration speed toward the scratched blank area and considerably reduced wound closure time. These results are consistent with WJMSCs-CM acting as a chemoattractant that drives directional fibroblast migration ^[20]^. Taken together, utilization of WJMSCs-CM successfully increases bio-instructive capability, effectively overcoming the stasis of HDFa healing in diabetic wounds and displaying remarkable skin tissue regeneration potential.

To further understand the mechanism by which WJMSCs-CM stimulated HDFa proliferation and migration, we examined the PI3K/Akt/eNOS pathway, a conventional central regulator of survival, metabolism, angiogenesis, and proliferation ^[21]^. WJMSCs-CM treatment significantly increased the protein expression levels of p-Akt and p-eNOS in HDFa, with activated p-Akt phosphorylating downstream eNOS at Ser1177 to generate its physiologically active form, p-eNOS ^[22]^. This finding is consistent with reports that restoring Akt phosphorylation in fibroblasts under hyperglycemic, diabetes-relevant conditions reactivates proliferation and migration alongside concurrent eNOS/VEGF upregulation and accelerated wound closure ^[23]^, supporting Akt/eNOS reactivation as a generalizable route by which diabetic fibroblast dysfunction can be pharmacologically regulated. In the early stages of wound healing, activation of p-eNOS in HDFa is critical for encouraging the local release of a low concentration of nitric oxide (NO) from cells. This endogenous NO molecule acts as a gas signaling molecule, promoting fibroblast skeleton reorganization and directed migration ^[24]^. The Akt/eNOS-NO pathway regulation, which is not confined to fibroblasts but also underlies the enhanced HUVEC angiogenic behavior ^[25]^, offers a plausible mechanistic bridge between the *in vitro* fibroblast/endothelial findings and the superior hair follicle regeneration observed *in vivo*. This is consistent with our histological findings, where the NF-AF dressing achieved the greatest follicle regeneration among the injured cohorts (∼17 follicles by day 21, versus <10 in untreated controls), suggesting the Akt/eNOS-driven activation captured *in vitro* is mechanistically continuous with the dermal appendage regeneration seen *in vivo*.

Angiogenesis is a complex physiological process involving re-establishing blood supply and reconstructing the microvascular network, which are similarly indispensable for overcoming the pathological ischemia and hypoxia that characterize diabetic chronic wounds ^[26]^. In this case, we evaluated the biological effectiveness of WJMSCs-CM efficiently protected HUVECs from high-glucose-induced damage and markedly enhanced their proliferation. This protective effect is consistent with reports that WJMSCs-CM restores viability, migration, and angiogenic capacity in high-glucose-damaged HUVECs through paracrine signaling ^[27]^, reinforcing that our TCM-primed WJMSCs-CM engages a conserved endothelial rescue pathway rather than a system-specific artifact. This benefit was more directly confirmed in tube formation assays with greater master segment length, node number, total length, master junctions, and branching length in CM-treated groups. Diabetic chronic wounds frequently exhibit a deficiency in pro-angiogenic signals, such as vascular endothelial growth factor (VEGF) and angiogenin, due to the specific requirements of this process ^[26]^. These results indicate that TCM stimulation converted WJMSCs into more effective producers of angiogenic factors. Combined with the fibroblast repair capacity described above, WJMSCs-CM therefore offers a dual biological repair strategy that simultaneously restores tissue proliferation and microcirculation, both of which are compromised in diabetic wound pathology. Having established the *in vitro* efficacy of WJMSCs-CM, we next addressed the challenge of delivering these paracrine factors to the wound bed within a controlled condition. We employed the coaxial eletrospun scaffold as a controlled-release vehicle for CM. TEM revealed a distinct core-shell fiber architecture, wherein the outer shell physically shields the WJMSCs-CM-loaded core from early degradation in the wound microenvironment ^[28]^.

At an optimized injection rate of 0.06 mL/h, SEM showed a uniform average fiber diameter of approximately 300 nm, closely mimicking the nanoscale dimensions of native ECM collagen fibers ^[29]^. This scale is favorable for promoting cell adhesion and migration through contact guidance. The scaffold’s high porosity further supports oxygen diffusion, nutrient transfer, and three-dimensional cellular and vascular infiltration, both of which are limiting factors in chronic diabetic wounds. Beyond its structural design, the scaffold’s material properties reinforced its translational suitability. A moderate swelling rate maintained wound moisture balance, while its progressive in vitro degradation profile was well synchronized with new tissue regeneration and ECM remodeling ^[30]^. The core-shell architecture also mitigated the burst release that commonly limits uniaxial scaffolds ^[31]^. The WJMSCs-CM within the core was released in a steady, progressive manner that avoided premature depletion of growth factors, sustaining molecular signaling across the prolonged healing timeline required for chronic wounds.

To ascertain the bioefficacy of the controlled-release scaffold in this investigation, cell viability demonstrated its superior cell compatibility. Moreover, the gradually released WJMSCs-CM from the core layer preserved its bioactivity, consistently delivering biochemical signals for proliferation. In relation to biomimetic topology, SEM analysis revealed remarkable cell dispersion on the nanofiber surface, with extensive pseudopodia and filamentous extensions creating an interconnected network that effectively emulates the spatial arrangement of the natural ECM. This morphological adaptation was corroborated at the molecular level; immunofluorescence labeling revealed markedly intensified signals and stress fibers in the cells of the experimental group. Exceedingly aligned with the activation of p-Akt/p-eNOS and the outcomes of scratch healing. In conclusion, our coaxial controlled-release scaffold effectively reestablished a three-dimensional, expansive, and directional migration via a dual approach of physical biomimetic topological cues and chemically regulated paracrine signals, showcasing remarkable potential for tissue engineering applications.

A transwell assay further showed that scaffold-released WJMSCs-CM established a chemotactic gradient sufficient to recruit HUVECs from the upper chamber, confirming that the delivery platform preserved the chemoattractant activity of the conditioned medium. In a multicellular co-culture system approximating the in vivo wound environment, ELISA showed elevated pro-collagen type I secretion alongside stable MMP-1 levels in fibroblasts, consistent with collagen accumulation rather than excessive degradation, together with upregulated VEGF and angiogenin in the HUVEC co-culture, corroborating the tube formation and viability improvements described above ^[32]^. Collectively, these protein-level data confirm that the coaxial scaffold functions as an effective delivery platform for the paracrine factors governing both ECM remodeling and angiogenesis. Because in vitro assays alone cannot capture the full complexity of chronic wound pathology, we next evaluated the scaffold in an STZ-induced diabetic rat wound model.

In this full-thickness diabetic wound model, the nanofiber dressing cohorts demonstrated accelerated healing, with the combined NF-AF group achieving the most complete repair. This outcome reflects not only the effect from WJMSCs-CM and the coaxial scaffold, but also the synergetic effect from AS-IV and FMN compounds. Beyond its role as a physical barrier against infection and moisture loss, the scaffold sustained the bioactivity of WJMSCs-CM at the wound site, extending the same p-Akt/p-eNOS-driven fibroblast signaling and VEGF/angiogenin-driven angiogenic cascade observed in vitro, thereby supporting pro-collagen accumulation, granulation tissue formation, and microcirculatory restoration in the ischemic wound bed. These outcomes indicate that the WJMSCs-CM-loaded coaxial dressing supports diabetic wound healing through multi-component, multi-stage regeneration rather than a single dominant mechanism.

To further validate the quality of skin regeneration from a histopathological perspective, H&E staining and quantitative analysis were performed. On day 21, the groups treated with NF-FMN and NF-AF dressings exhibited an epidermal thickness most comparable to intact skin, confirming that the coaxial structure effectively guided orderly epithelial cell arrangement and enhanced mature re-epithelialization. Notably, a marked increase in newly formed hair follicles was observed in these treatment groups, indicating that sustained WJMSCs-CM release successfully stimulated endogenous hair follicle stem cell niches. This observation is consistent with the results of our *in vitro* study, which demonstrated enhanced hair follicle regeneration, accelerated healing, and increased angiogenesis in a diabetic mouse wound model ^[33]^. Furthermore, quantitative Masson’s trichrome analysis confirmed that the combined NF-AF treatment yielded significantly higher collagen density in late-stage repair, displaying a parallel, tightly packed wavy pattern characteristic of healthy dermis and aligning with our *in vitro* ELISA data. This optimal collagen remodeling stems from the sequential, controlled delivery of multiple bioactive agents, early released NF-FMN rapidly restores microcirculation, while sustained NF-AS-IV release continuously drives fibroblast collagen synthesis and assembly, counteracting the chronic collagen deficit in diabetic wounds. Consequently, these combined H&E and Masson’s outcomes demonstrate that the coaxial dressing effectively directs stage-specific tissue regeneration *in vivo*, achieving dual structural and functional repair.

To characterize the *in vivo* anti-inflammatory and immunomodulatory mechanisms underlying these outcomes, we performed IHC staining of wound tissue following the various treatments. The expression of pro-inflammatory M1 macrophage markers, CD68 and iNOS, and the inflammatory mediator COX-2, was significantly downregulated, while the anti-inflammatory M2 marker CD206 was markedly upregulated. These findings suggest that the dressing drove M1-to-M2 macrophage polarization and re-established a pro-healing microenvironment. The myofibroblast marker α-SMA exhibited distinct temporal dynamics consistent with the histological results. An early peak at day 7 in the NF-AF group supported rapid initiation of granulation tissue formation, in contrast to the delayed peak observed in control and comfeel-treated wounds at day 14. Furthermore, α-SMA expression subsequently resolved toward baseline by day 21, limiting the risk of excessive fibrosis and hypertrophic scarring ^[34]^. Taken together with the H&E and Masson’s trichrome findings, these results indicate that the TCM-functionalized coaxial dressing orchestrates diabetic wound regeneration in vivo through a coordinated, multi-dimensional mechanism spanning immunomodulation, angiogenesis, and ECM remodeling.

Several limitations should be considered when interpreting these findings. As a complex biological product, WJMSCs-CM is subject to batch-to-batch variability, and its precise active constituents remain to be fully defined. All *in vivo* comparisons in this study were made against an untreated control and a single commercial dressing (Comfeel), so the relative performance of this platform against other active wound-care technologies remains untested. The STZ-induced rat model, while standard for diabetic wound research, is a small-animal, short-term (21-day) system that does not capture the larger wound dimensions, slower healing kinetics, or comorbidities typical of human diabetic ulcers, and the long-term release kinetics and bioactivity retention of WJMSCs-CM beyond this window remain unknown.

Future work should extend evaluation to larger-animal models and longer healing timelines to better approximate clinical translation. Taken together, this study demonstrates that TCM-preconditioned WJMSCs-CM, delivered through a coaxial electrospun scaffold, restores fibroblast and endothelial function via Akt/eNOS signaling in vitro and translates this rescue into accelerated wound closure, immunomodulation, and hair follicle regeneration in a diabetic rat model, providing a mechanistically grounded foundation for further preclinical development.

## Conclusion

In this study, we successfully fabricated a novel electrospun nanofiber dressing characterized by a core-shell architecture and biomimetic physical properties, establishing a highly effective biomaterial platform for the dual and sustained delivery of herbal-primed stem cell secretomes. The design is developed to overcome the primary clinical limitations of conventional protein therapies, namely rapid enzymatic degradation and uncontrolled burst release in the protease-rich diabetic wound bed. *In vitro*, preconditioning with AS-IV and FMN significantly enhanced the paracrine function of WJMSCs-CM, promoting cell viability and activating the p-Akt/p-eNOS signaling pathway in HDFa, while stimulating three-dimensional transmembrane migration and capillary-like tube formation in HUVECs. Concurrently, these bioactive factors regulated ECM remodeling in co-culture models, suppressing excessive MMP-1 degradation while upregulating the secretion of pro-collagen type I, VEGF, and angiogenin to establish a molecular foundation for tissue regeneration and microvascular reconstruction. *In vivo*, the NF-AF dressing exhibited superior synergistic healing efficacy in an STZ-induced diabetic rat model. Histopathological and immunohistochemical analyses confirmed that this dressing modulated the immune microenvironment by driving M1-to-M2 macrophage polarization (downregulating CD68, iNOS, and COX-2 while upregulating CD206) and facilitating healthy wound contraction through dynamic ⍺-SMA regulation. By day 21, the system achieved physiological restoration of epidermal thickness, orderly mature collagen deposition, and regeneration of functional hair follicle appendages. Crucially, this study demonstrates a profound synergistic effect achieved by co-delivering secretomes primed by two distinct herbal compounds, Astragaloside IV and Formononetin. Evaluated in a severe Type 1 diabetic rat model, this dual-functional scaffold orchestrates the complete tissue repair cascade, transitioning the hostile wound microenvironment from a chronic pro-inflammatory state to an active remodeling phase, ultimately achieving a remarkable 98.2% wound closure that mathematically outpaces the additive effects of single-agent therapies. Overall, the coaxial controlled-release nanofiber dressing integrates biomimetic topological cues with biochemical signaling, highlighting its strong translational potential for diabetic skin tissue engineering

## Authorship contribution statement

**Kuo-Hui Chiu**: Writing – original draft, Investigation, Formal analysis. **Lin-Chu Huang**: Writing – review & editing, Validation, Conceptualization. **Wen-Ling Wang**: Writing – review & editing, Conceptualization, Project administration. **Yi-Hui Lai**: Methodology, Data curation. **Chun-Hsu Yao**: Supervision, Funding acquisition, Conceptualization.

## Declaration of competing interest

The authors declare that they have no known competing financial interests or personal relationships that could have appeared to influence the work reported in this paper.

## Declaration of generative AI and AI-assisted technologies in the manuscript preparation process

During the preparation of this work the authors used Claude in order to language editing. After using this tool, the authors reviewed and edited the content as needed and take full responsibility for the content of the published article.

## Acknowledgments

This study was supported by grants CMU113-MF-60 and CMU114-S-03 from China Medical University. Figure 4A and figure 7A were created with BioRender.com.

## Notes

### Competing Interest Statement

The authors have declared no competing interest.

